# A Markovian neural barcode representing mesoscale cortical spatiotemporal dynamics

**DOI:** 10.1101/2024.06.29.601346

**Authors:** Jordan M Culp, Donovan M Ashby, Antis G George, G. Campbell Teskey, Wilten Nicola, Alexander McGirr

## Abstract

Mesoscale cortical dynamics consist of stereotyped patterns of recurring activity motifs, however the constraints and rules governing how these motifs assemble over time is not known. Here we propose a Continuous Time Markov Chain model that probabilistically describes the temporal sequence of activity motifs using Markov Elements derived using semi-binary non-negative matrix factorization. Although derived from a discovery sample, these can be applied to new recordings from new mice. Unwrapping the associated transition probability matrix creates a ‘Markovian neural barcode’ describing the probability of Markov element transitions as a compact and interpretable representation of neocortical dynamics. We show broad utility across a range of common mesoscale cortical imaging applications, ranging from time-locked events to pathological models. Moreover, it allows the discovery of new and emergent Markov Elements that unmask the flexibility of constraints governing cortical dynamics. The Markovian neural barcode provides a novel and powerful tool to characterize cortical function.

## Introduction

Brain activity is highly structured and reflective of the behavioral state of the animal^1^. Mesoscale cortical imaging samples this structured activity from a large expanse of dorsal neocortex simultaneously^2^ and can leverage haemodynamic signals, organic dyes or an increasingly diverse set of genetically encoded indicators with cell-type specific expression^3-6^. The versatility of the approach has led to its application to a breadth of questions in neuroscience, ranging from sensory processing^7-9^, neurotransmitter systems^3^, learning and memory^10,11^, behavior^12,13^, decision making^14^, aging^15^, and disease models^16-18^.

Across applications, mesoscale cortical imaging involves sampling neural signals that are recurring, stereotyped and involve anatomically and functionally distinct regions of neocortex^19-22^. In some applications, these activity motifs introduce variability into the time locked signal of interest, for example a sensory response^7-9^ or a behavioral response in the absence of a cue^14^. In others, these activity motifs are not spontaneous at all, and reflect a direct product of the animal’s behavior or state^12,23^. Alternatively, some mesoscale imaging applications remain agnostic to the cause of activity motifs, and instead leverage the sequence of motifs as the basis for functional connectivity to define large scale interaction networks^24-28^. Although there is ongoing debate as to the role and function of these recurring activity motifs^29,30^, their preservation at significant energetic and wiring cost belies their importance^31-33^. Moreover, the means by which motifs of neural activity assemble over time and the rules governing these transitions provides a novel lens from which to view the structure of cortical dynamics^34^ as the fundamental architecture of large scale networks^35,36^.

Previous approaches to resolving this complexity in mesoscale cortical imaging have included unsupervised methods of motif detection^34^ or experimentally generated templates of neural activity^19,20,26,37-39^. While elegant solutions, existing unsupervised approaches require computational resources that limit scalability and experimental templates are best suited to circumscribed neural circuits. Unsupervised methods that are generalizable across applications of mesoscale imaging, and other high-dimensional sampling methods, require a compact lower-dimensional representation of dynamics and their constraints^34,40^.

Here, we present a method and formal model for doing so that we term the ‘Markovian neural barcode’. This approach discretizes mesoscale cortical dynamics with a generalizable set of cortical activity motifs. We then utilize a Continuous Time Markov Chain (CTMC) model^41^ to extract the temporal structure of mesoscale cortical activity motifs. The probabilistic sequence of these Markov Elements reveals constraints governing neocortical dynamics that can be visualized as a ‘Markovian neural barcode’.

## Results

### Mesoscale Imaging Reveals Markovian Rules Governing Cortical Dynamics

We performed *in vivo* mesoscale cortical imaging on a total of 123 (55 Females and 68 Males) awake head fixed mice (**Figure 1A**). Transgenic Thy1-jRGECO1a mice^42^ expressing the red shifted calcium indicator received chronic windows exposing dorsal neocortex, with registration to the Allen Brain Atlas and application of a mask to remove edge artifacts as well as the sagittal sinus and its major tributaries (**Figure 1B**). The cortical dynamics sampled with mesoscale cortical imaging are shown in montages at different timescales in **Figure 1C** to illustrate the diversity of spatiotemporal activity in neocortex.

**Figure 1.**
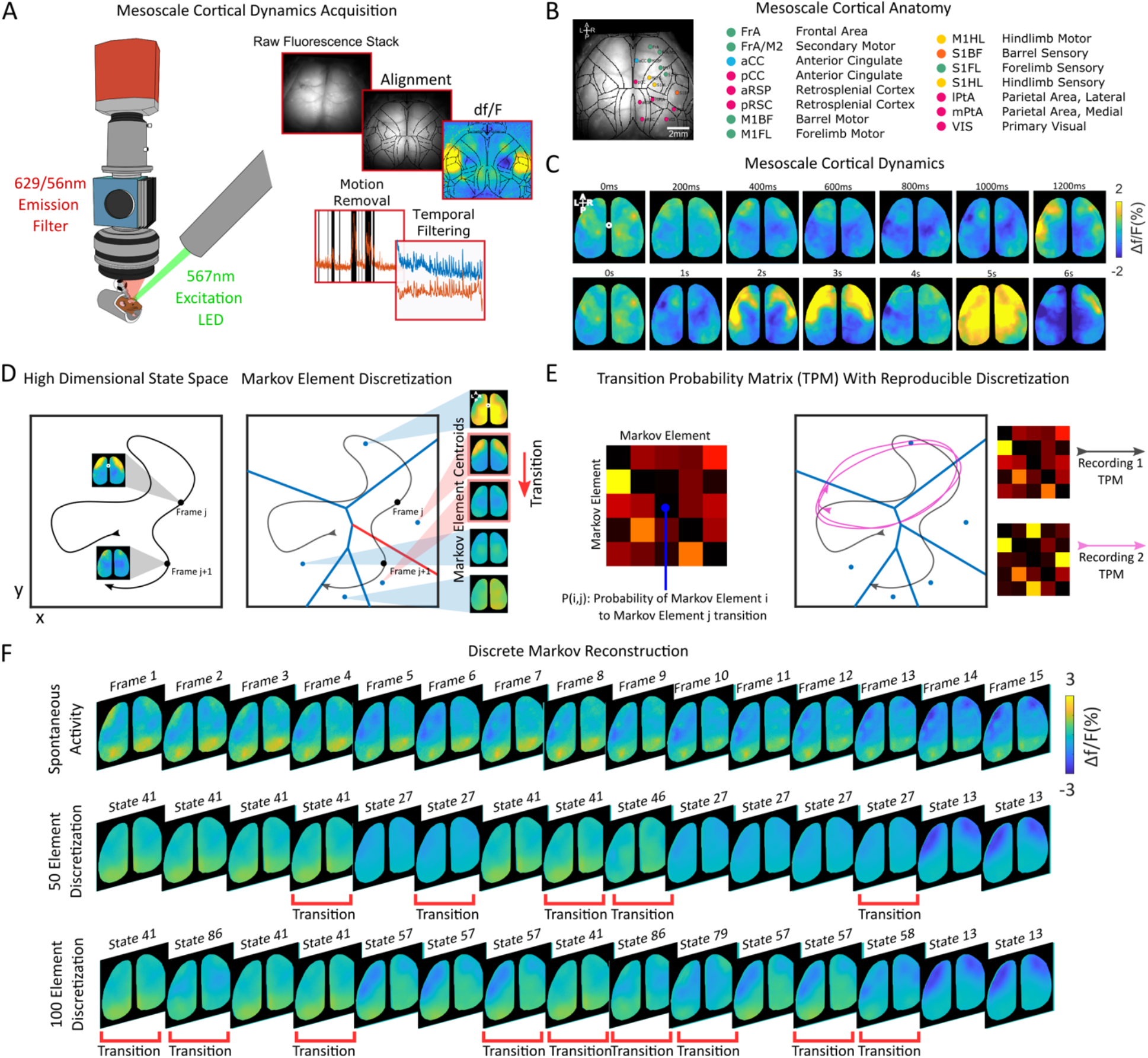
Markovian dynamics in mesoscale cortical calcium activity. **(A)** A schematic representation of mesoscale cortical imaging and pre-processing of the recordings. The raw fluorescence stack is first aligned to the Allen Institute Mouse Brain Atlas, followed by spatial filtering to identify epochs of movement with sensitivity comparable to behavioral cameras. Following temporal filtering the final signal Δ*F*/*F* is computed. **(B)** An annotated mesoscale fluorescence image is shown after alignment to the Allen Institute Mouse Brain Atlas. **(C)** In the upper montage, mesoscale cortical imaging spatiotemporal dynamics are illustrated with frames every 200ms, and in the lower montage are frames selected every 1s. **(D)** A schematic representation of discretizing high dimensional state space into a series of Voronoi cells in a Voronoi diagram, where each cell is described by the centroid or Markov element. The trajectory of frames from mesoscale cortical imaging can be approximated by the Markov element with the transitioning between centroids modeled as a CTMC. **(E)** Schematic representation of (Left) the transition probability matrix estimated with the relative frequency of transitioning from one Voronoi cell to another. The entry *P*(*i, j*) in this matrix denotes the probability of moving from state *i* to state *j*, where the states correspond to the Voronoi cells in Figure (D). (Right) The grey system is a schematic of a chaotic regime, whereas the pink system is a schematic of a more oscillatory dynamical regime with its associated sparser transition probability matrix. **(F)** The application of Markov element assignment with increasing number of Markov Elements for a CTMC model. Note the relative paucity of transitions with fewer Markov elements, and as more Markov elements are used the approximation the CTMC approximation to the mesoscale imaging data also improves.

The “raw” calcium dynamics spans a high-dimensional state-space consisting of potentially thousands of independent pixel trajectories (**Figure 1C; SVideo 1**). However the overall dimension of the raw dynamics was low (**SFigure 1**), indicating that individual pixels of calcium activity trajectories are not independent in high dimensional space^23^ and can occupy a low dimensional subspace. Accordingly, though the mesoscale recordings suffer from a conventional ‘big-data’ problem, the low-dimensional dynamics allow us to consider a more compact dynamical system.

Thus, we applied a novel, state-space discretization approach conceptually illustrated in **Figure 1D&E**. The state space can be discretized into ‘zones’ each defined by a central Markov Element that is the high dimensional space centroid determined using unsupervised semi-binary non-negative matrix factorization (SBNNMF)^43,44^. Note that alternative clustering algorithms (e.g. k-means) can also create a suitable set of Markov Elements, however require expert curation. The high-dimensional calcium dynamics can then be tracked as crossings from one ‘zone’ to the next (**Figure 1E**). This then informs a Transition Probability Matrix (TPM) whose elements *TPM* (*i, j*) yield the probability of crossing from zone *i* to zone *j* (**Figure 1E**). If generalizable constraints govern the transitions between zones, then the transition probability matrices that emerge will be reproducible even if activity is predominantly cycling between different zones (i.e. the recording’s Markov Element ‘occupancy’ distribution is different). Cycling with a different trajectory will produce different TPMs representing a deviation from the generalized rules governing Markov Element transitions, whether the result of a normative, experimental, or pathological process (**Figure 1E**).

To determine the robustness of the Markovian neural barcode, we first tested the reproducibility of Markov Elements derived across mice. From a discovery sample consisting of 40 frames per recording randomly selected from 80 recordings from 20 mice (the 3,200 frames discovery sample), the SBNNMF algorithm identified 140 Markov Elements. Ranked according to the number of frames to which they were assigned, 95% of the discovery sample was explained by the first 100 Markov Elements (**SFigure 2A**). To validate the set of Markov Elements, we repeated the SBNNMF algorithm with 40 frames per recording drawn from an independent set of 80 recordings from 20 mice (the 3,200 frame validation sample). The ‘discovery’ and ‘validation’ set of Markov Elements was a highly consistent as were rank assignments (**SFigure 2B**). To further cross-validate, we then derived two new sets of Markov Elements from the ‘discovery’ recordings by subsampling 40 frames (Random Validation Basis Set 1) and 80 frames (Random Validation Basis Set 2) from each recording, which also revealed highly consistent Markov Elements and highly correlated rank assignment (**SFigure 2C**). At its extreme, we asked whether data from a single recording from a single mouse would provide a set of Markov Elements consistent with the discovery sample. This also revealed a high degree of concordance and rank correlation with the ‘discovery’ set of Markov Elements (**SFigure 2D**). Therefore, the Markov Elements derived from one initialization of SBNNMF can be generalized to novel acquisitions, and importantly this occurs without an excess of ‘high reconstruction error frames’, defined as >3SD Euclidean difference between the raw frame and the assigned Markov Element (**SFigure 2E**).

A potential concern when discretizing a high dimensional dynamical system is the artificial introduction of ‘transitions’ when the raw data is at the boundary between two centroids. This can be minimized by adding a local (in time) penalty on transitions. This cost function balances minimizing the reconstruction error with the total number of transitions (**SFigure 2F**), resulting in a sparser yet still highly correlated transition probability matrix and continuous Markov chain reconstruction (**SFigure 2G&H**). After this optimization, one can determine the reconstruction error associated with increasingly fine discretization with the discovery basis set (**SFigure 3**), and model the number of frames between transitions, or conversely the ‘dwell’ or ‘holding’ time in each Markov Element from the continuous time Markov chain (**SFigure 4**). Our data suggests that mesoscale cortical dynamics can be adequately approximated and quantified with a finite number of Markov Elements, recognizing that a full reconstruction of high-dimensional dynamics would require infinite elements^45-47^.

A montage illustrating the discrete Markov reconstruction is shown in **Figure 1F**, with the raw recording in the top panel and finer grained reconstructions below and also depicted in **SVideo 2**. Critically, the CTMC description is a formal generative model of brain dynamics, with novel, synthetic, realizations of any recording easily generated with the holding times and the transition probability matrices (**SVideo 3**).

To determine whether there is structure to Transition Probability matrices, we applied Louvain community detection to reveal four modules within the transition probability matrices (**Figure 2A**). The functional organization of these modules indicate that modules serve as pseudo-’absorbing states’ in a Markov process, with prominent intra-module transition blocks. To facilitate visualization, the Louvain sorted average transition probability matrix is presented as a color-coded transition probability matrix making intra- and inter-modular transitions between Markov Elements readily apparent.

**Figure 2.**
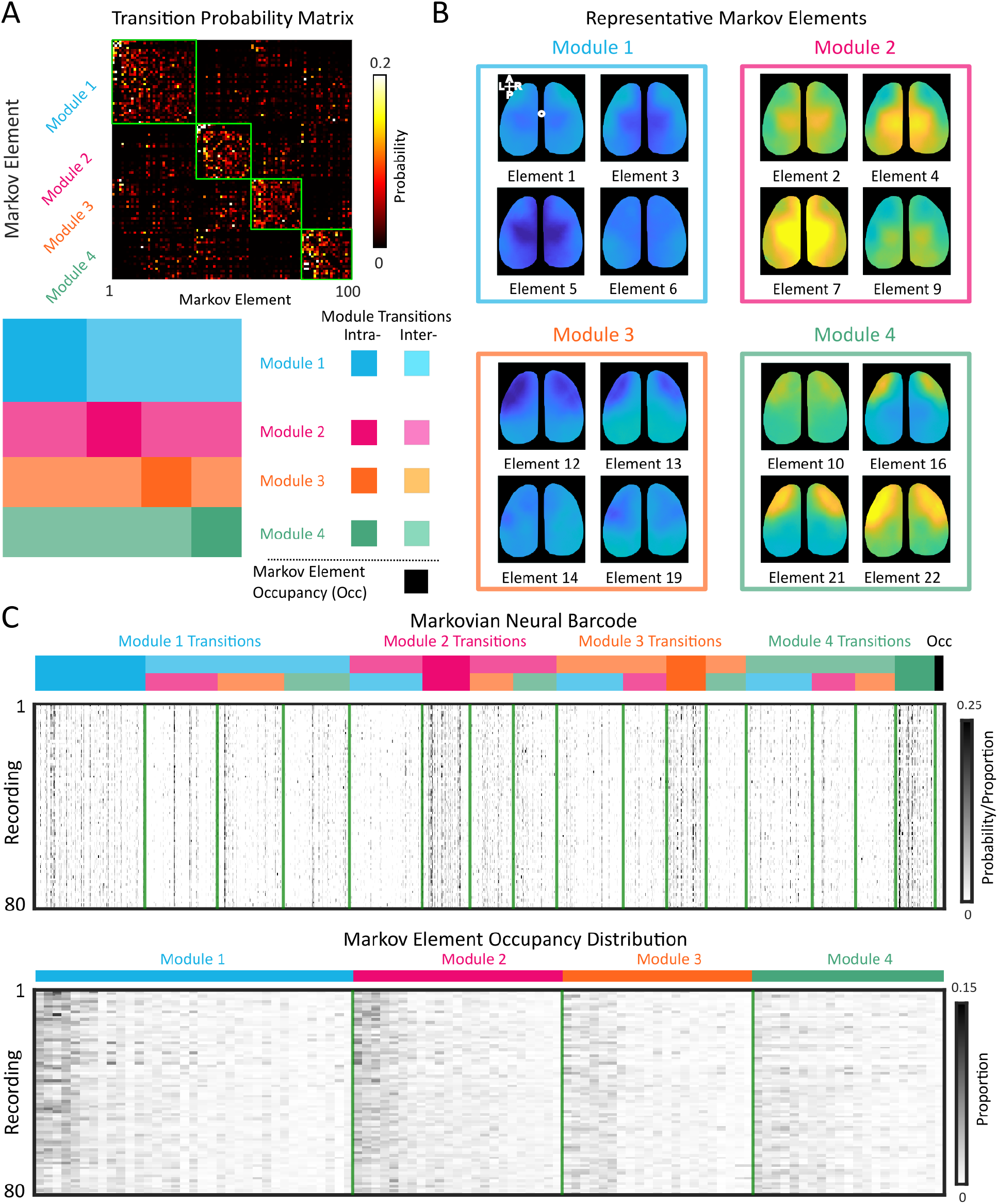
The Markovian neural barcode. **(A)** (Top) A Louvain sorted transition probability matrix for a single recording with boundaries between the modules denoted by a green border. (Bottom) A schematic of the transition probability matrix is shown color coded for transitions and the Markov element occupancy distribution. **(B)** Representative color-coded Markov Elements depicted after Louvain community detection applied to transition probability matrices. **(C)** The Markovian neural barcode. (Top) A schematic color-coded legend to facilitate visualization and interpretation of the barcode and individual rows for n=80 recordings. (Bottom) A schematic color-coded legend of the occupancy distribution. All entries in the neural barcode are probabilities, with darker colours denoting higher probabilities.

Representative Markov Elements from each of these modules are illustrated in **Figure 2B**. The Markov Elements from each module were distinct but shared some general characteristics along anterio-posterior and medio-lateral gradients that have previously been described^34,35,38,48^. Note that Markov Elements from Module 1 show a medial-lateral ΔF/F gradient in which midline and primary somatosensory regions have lower ΔF/F signal compared to lateral supplementary sensory regions, anterior and posterior cortical regions. For Markov Elements in Module 2, calcium signal is prominent in midline cortical regions including the cingulate, the retrosplenial cortex, parietal association area and supplementary motor regions. Modules 3 and 4 were distinguished by their anterio-posterior ΔF/F signal gradients, with Module 3 having greater ΔF/F signal in visual areas and lower ΔF/F signal in facial and barrel motor regions, where as Module 4 was typified by the converse.

The Louvain sorted transition probability matrix and its associated color-coded legend forms the basis of the legend for the unwrapped transition probability matrix illustrated in **Figure 2C**, which we term the ‘Markovian neural barcode’. The lower row of the legend denotes the Markov Element of origin and its associated color-coded module, and the upper row denotes the Markov Element that has been transitioned to and its associated color-coded module. In the Markovian neural barcode, each row corresponds to the unwrapped transition probability matrix derived from a single recording. Also depicted is a zoom of the Markov Element occupancy distribution, denoted by the black bar in the barcode legend.

An important feature of the Markovian neural barcode is its tractability for further quantification and visualization. Indeed, three principal components explained 73% of the variance (**SFigure 5A&B**). Accordingly, principal component projections of the Markovian neural barcode according to these first three components provides visualization of clustering, and statistical quantification of intra- and inter-distances. Though we favor principal component analysis, other dimensionality reduction techniques can also be applied to the Markovian neural barcode (**SFigure 6**).

### Acquisition Parameters and the Markovian Neural Barcode

To determine whether the Markovian neural barcode was robust to different acquisition parameters, we varied the shutter speed to acquire mesoscale cortical activity at 10Hz (99.7ms), 30Hz (33.1ms), and 50Hz (19.7ms). This creates variation in the spatiotemporal composition of the raw calcium signal to test the accuracy of the continuous Markov time chain reconstruction using Markov elements (acquired at 50Hz). As expected, faster acquisition rates were associated with more transitions for recordings of identical duration (**SFigure 7A&B**), however the reconstruction error did not increase with different sampling rates (**SFigure 7C&D**). The composition of Markovian neural barcode was highly conserved across different shutter speeds, however dynamical differences emerge as a function of temporal averaging with longer exposure (**SFigure 7E-H**).

An important consideration when using blue-green indicators is the impact of haemodynamic artifacts on the fluorescence signal^49^, and potentially the associated Markov Element to which a frame is assigned. Red-shifted indicators, such as jRGECO1a, are less susceptible but not immune to these confounds^28,50^. We found no statistically significant differences in the Markovian neural barcode comparing haemodynamically corrected^50^ and uncorrected signals for *n*=22 recordings (**SFigure 8A-C**). Moreover, the impact of haemodynamic correction on intra-recording distances was small (**SFigure 8D**).

### Validation of the Mesoscale Cortical Imaging Markovian Neural Barcode

To determine whether continuous trajectories can be described with our approach, we considered the Lorenz system of ordinary differential equations defining a classical chaotic system (**SFigure 9**) where attractors are generated by two slightly different parameter sets (**SVideo 4**). These attractors are depicted along their x, y, and z axes (**SFigure 9A**) and as a two-dimensional representation in phase space from which Markov Elements can be extracted (**SFigure 9B**). From this simulation, the Markov Elements explaining 95% of the ‘frames’ (**SFigure 9C**) reveal a transition probability matrix with Louvain sorted modules for stereotyped transitions within and between modules (**SFigure 9D**) and their associated Markovian barcode for each set of attractors (**SFigure 9E-H**). Accordingly, discretize reconstruction of continuous trajectories is possible as a continuous Markov time chain.

Next, we tested the Markovity of mesoscale cortical dynamics and their continuous Markov time chain reconstructions. For these analyses, we used Bayesian Information Criterion (BIC) as a method to determine model fit among a second-order Markov process relative to a first-order, and a zeroth-order process^51^. We seek a model that achieves the optimal balance between goodness of fit with a parsimoniousness selection of model parameters, and therefore compare zeroth- and first-order processes to a (higher) second-order process^51-54^. This models independence of future state from current state (stochastic zero-order), as compared probability based the current motif (first-order), and doublet sequences (second-order). Our data suggests that mesoscale cortical calcium imaging data is best described as a first-order Markov process (**SFigure 10**).

We next sought to determine the internal consistency and generalizability of the Markovian neural barcode. We first asked how much data is required for the Markovian neural barcode to converge. This revealed that the transition probability matrices converge within approximately 200 seconds of data acquired at 50Hz (**SFigure 11A**). We next examined the generalizability of the transition probability matrices are across animals. Across the *n*=80 recordings in the ‘discovery’ sample, transition probability matrices are very reproducible with r=0.70. To determine the chance occurrence of such reproducibility, we used two approaches wherein the original recording occupancy distributions were preserved (**SFigure 11B**). First, we shuffled the continuous chain Markov Element assignment logs and found that recordings were correlated at 0.19. Then, we shuffled the rows of the transition probability matrices and found no correlation between recordings (r=-0.002). Finally, we examined the generalizability of Markov Elements occupancy distributions giving rise to these highly consistent transition probability matrices across recordings (**SFigure 11C**), and found that these were much more variably conserved across recordings (r=0.56), however that multiple recordings from the same animal (*n*=80) showed very reliable occupancy distributions (r=0.85).

Accordingly, a first-order Markov process reliably describes mesoscale cortical dynamics, converging with relatively short acquisitions, to generate transition probability matrices that are generalizable across animals.

### The Markovian Neural Barcode and Sensory Evoked Responses

Transition probabilities from a continuous time Markov chain may be homogeneous, or it may become inhomogeneous when an animal is engaging with the environment. Thus, the trial-by-trial dynamics following a time-locked event such as a sensory stimulus can inform whether pre-existing activity trajectories continue despite sensory evoked stimuli, with important implications for how neural states stage appropriate responses to specific contexts. We therefore examined sensory evoked responses resulting from full field flashes directed to the right eye (5ms, 450nm LED, Thorlabs; **Figure 3A**). The LED illuminated every 3 seconds with a 1 second jitter to minimize anticipation and habituation, without any task or reward contingencies.

**Figure 3.**
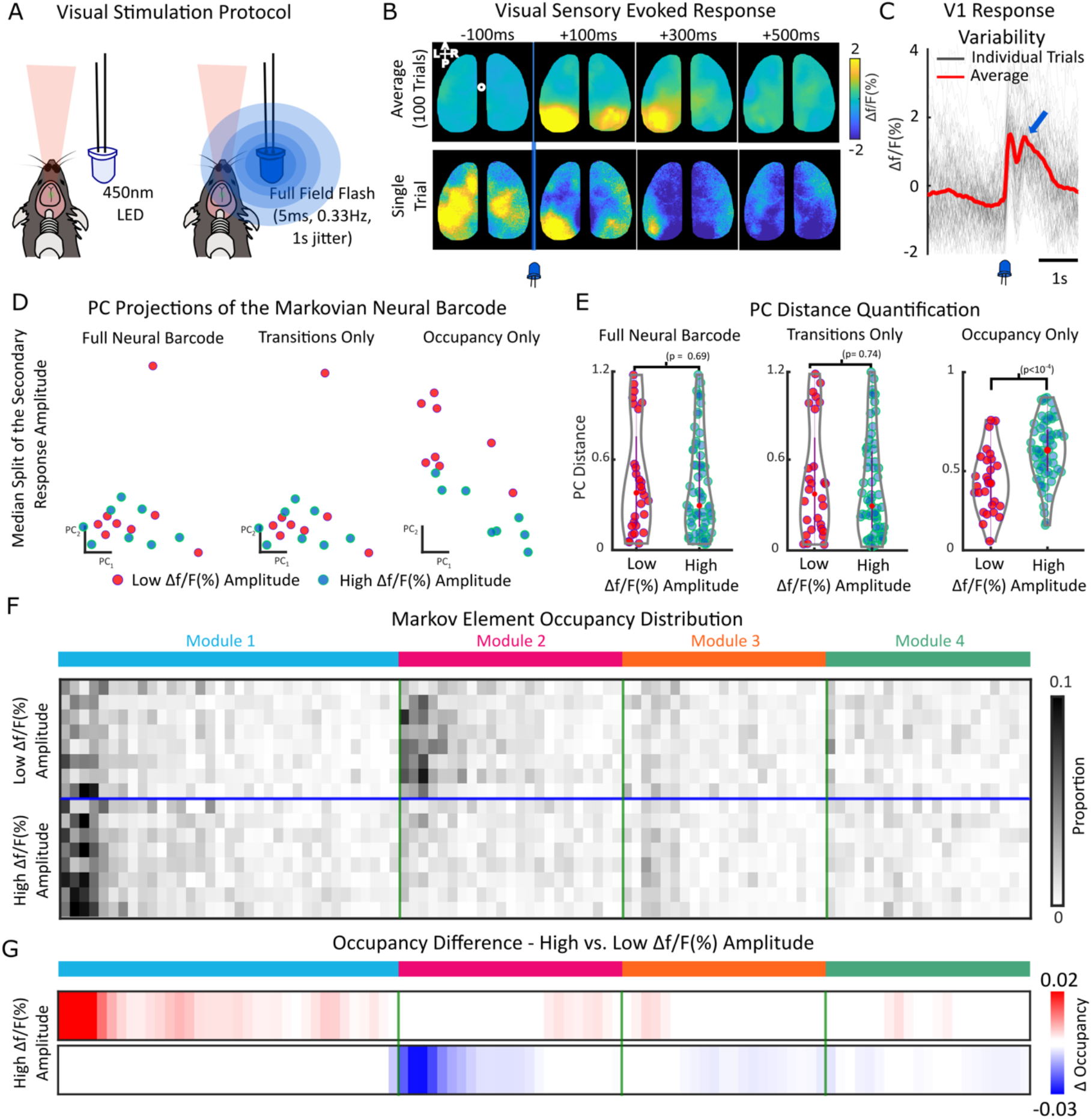
Sensory evoked processing is evident in the Markovian neural barcode. **(A)** A schematic representation of mesoscale cortical imaging during full field visual flash experiments. **(B)** A “raster plot” of the Markov State occupancy (y-axis) vs time (x-axis), with the states colour coded according to the Louvain sorted activity modules. **(C)** The histogram of the Markov State occupancy for the second preceding and following a full field LED flash. **(D)** The Louvain sorted transition probability matrices derived from transitions occurring prior to the full field visual flash (Left) and after the flash onset (Right). Note the upregulation of inter-module transition probabilities. **(E)** The Markovian neural barcode for the same animals imaged under a control quiet wakefulness condition and with field visual flashes. **(F)** To visualize and facilitate interpretation, we display the mean difference in the Markovian neural barcode between visual stimuli and control recording for transition probabilities and occupancy distribution. **(G)** The first two principal component projections of the neural barcode for the full barcode (left), the transition probability matrices only (middle), and the occupancy distribution only (right) for the control (white circle) and visual stimulation protocol recordings (blue circle). **(H)** Statistical quantification of the PC distance reveals statistical separation for the full neural barcode ((Left) Wilcoxon Rank Sum test; Control: 0.2517,Visual: 0.5045, *p* = 3.1908*e* − 3), the transition probability matrices only ((Middle) Wilcoxon Rank Sum test ; Control: 0.2507, Visual: 0.5041, *p* = 3.2795*e* − 03) and the occupancy distribution only ((Right) Wilcoxon Rank Sum Test; Control: 0.3670, Visual: 0.5829 *p* = 4.9088*e* − 3)

To visualize the impact of sensory evoked responses we plotted the time-series of Markov Elements as an occupancy raster (**Figure 3B**). The impact of visual stimulation was apparent as a trial-aligned average within the first 500ms (25 frames) following stimulation (**Figure 3C**). It is important to note that upregulated Markov Elements belong to all four Modules (**Figure 3C**). To examine how visual stimuli alter transition probabilities, we illustrate transition probability matrices derived from the 500ms preceding and following a visual stimulus (**Figure 3D**), revealing an upregulation of inter-module transitions following visual stimuli. To statistically quantify this without the confounds of visual stimulus carryover, we acquired two samples from each mouse, first under a control condition in the absence of visual stimuli, and second while the animal received visual stimuli as in **Figure 3A**. The Markovian neural barcode for these conditions is illustrated in **Figure 3E**, and the difference between conditions visualized in **Figure 3F**. Using PC projections of the barcode reveals clustering and separation of visual stimulus recordings and control recordings (**Figure 3G**), as confirmed by quantification of the PC distance distributions (**Figure 3H**). The constraints governing cortical activity motif transitions therefore reflect both homogeneous and inhomogeneous processes, where transition probabilities represent an integration of sensory information, internal state, and spontaneous activity^23,31,55^. Accordingly, appropriate action selection requires flexibility in the constraints governing cortical dynamics as revealed by the Markovian neural barcode.

The visual stimulation produced a consistent visual neocortical response evident in the 100-trial average, however the individual trials varied considerably as visual responses overlay ongoing spontaneous cortical dynamics^20,56^ (**Figure 4A-C; SVideo 5**). This variability in response amplitude was most pronounced for the secondary component of the response (**Figure 4C**).

**Figure 4.**
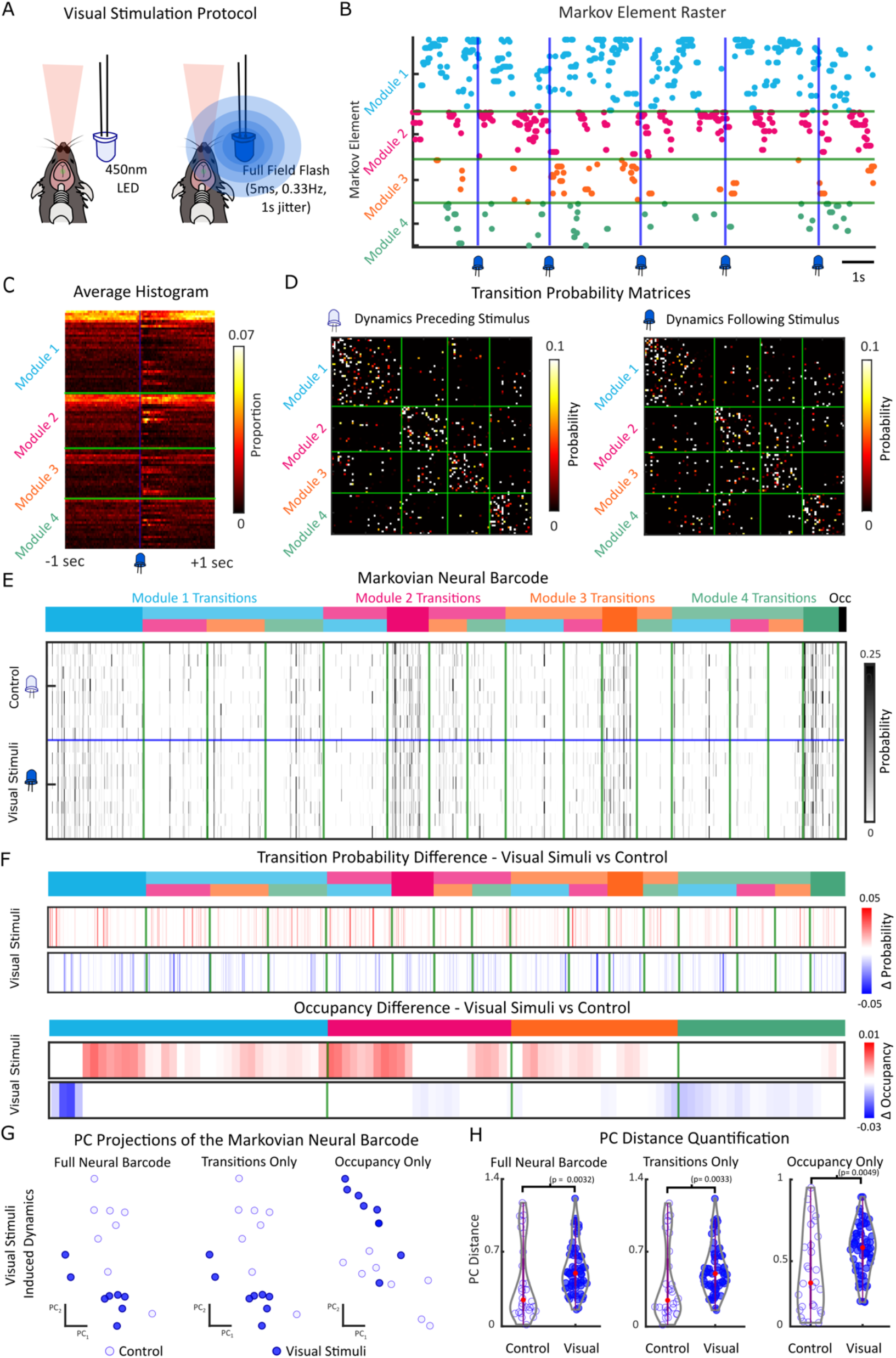
The Markovian neural barcode explains variability in sensory evoked processing. **(A)** A schematic representation of mesoscale cortical imaging during full field visual flash experiments. **(B)** Visual evoked responses from a full field flash produce a characteristic response in V1 (top: 100-trial average; bottom: single-trial). Note the considerable variability in the late component (2300ms) of the visual evoked response. **(C)** The heterogeneity in visual evoked response amplitudes is shown on a per trial basis. The average visual evoked response trace is shown in red, revealing the primary and secondary visual evoked responses. **(D)** Projections of the Markovian neural barcode for visual flash trials dichotomized according to a median split of the secondary response amplitude from (n=8) mice. **(E)** Quantification of the PC projections from the first three components reveals that only the Markov element occupancy distribution distinguishes visual evoked response trials with high and low Δ*F*/*F* secondary responses (Wilcoxon Rank Sum test; (Full) Low: 0.3870, High: 0.3004, *p* = 6.9315*e* − 01; (TPM) Low: 0.3822, High: 0.3024, *p* = 7.3749*e* − 01; (OCC) Low: 0.3766, High: 0.5639, *p* = 6.5393*e* − 05). **(F)** The Markov element occupancy bar code is presented for n=8 animals for trials that elicited high and low Δ*F*/*F* secondary responses. Note upregulation of Module 2 Markov elements in low Δ*F*/*F* amplitude secondary response trials, and an upregulation of Module 1 Markov elements in high Δ*F*/*F* amplitude secondary response trials. **(G)** To better visualize the difference between conditions, we express the Markovian neural barcode as the absolute difference between the proportion of each Markov element occupancy.

In mice and humans, the amplitude of this secondary visual evoked response has been associated with enhanced visual discrimination for subsequent stimuli ^57^. We therefore examined lower dimensional PC-projections of the Markovian neural barcode as a function of a median split of the baseline corrected secondary visual evoked response amplitude (**Figure 4D**) with data limited to the second preceding each visual stimulus. The PC-projections revealed no clustering for the full Markovian neural barcode or the transitions that dominate it, however there was evidence of clustering in the Markov Element occupancy distribution, as confirmed with PC distance quantification (**Figure 4E**). The occupancy distribution is illustrated in **Figure 4F**, in which low ΔF/F secondary response amplitude trials have a clear overrepresentation of Module 2 Markov elements preceding the stimulus as opposed to an overrepresentation of Module 1 Markov elements in high ΔF/F trials, most clearly visualized as a difference barcode revealing up and downregulated Markov Elements (**Figure 4G**). This is interesting, because the regional composition of activity in Module 2 elements has been linked to the murine equivalent of the ‘default mode network’, a network that must deactivate for successful task performance^58^.

### Dynamical Signatures are Impacted by Extreme Excitation

Next, we used maximal electroconvulsive seizures (MES) (**Figure 5A&B**) to ask how the constraints governing cortical dynamics would be altered following extreme excitation^59^ as an exemplar of pathology. Seizures are a powerful test of brain dynamics because the induced hypersynchronous and hyperexcitable neural events results in a predictable pattern of electrical activity, whereby the seizure is followed by suppression of activity dynamics, and then a gradual increase in neural activity power (**SVideo 6**). Nevertheless, physiology^60^ and behavior^61^ remain persistently disrupted after termination of the seizure.

**Figure 5.**
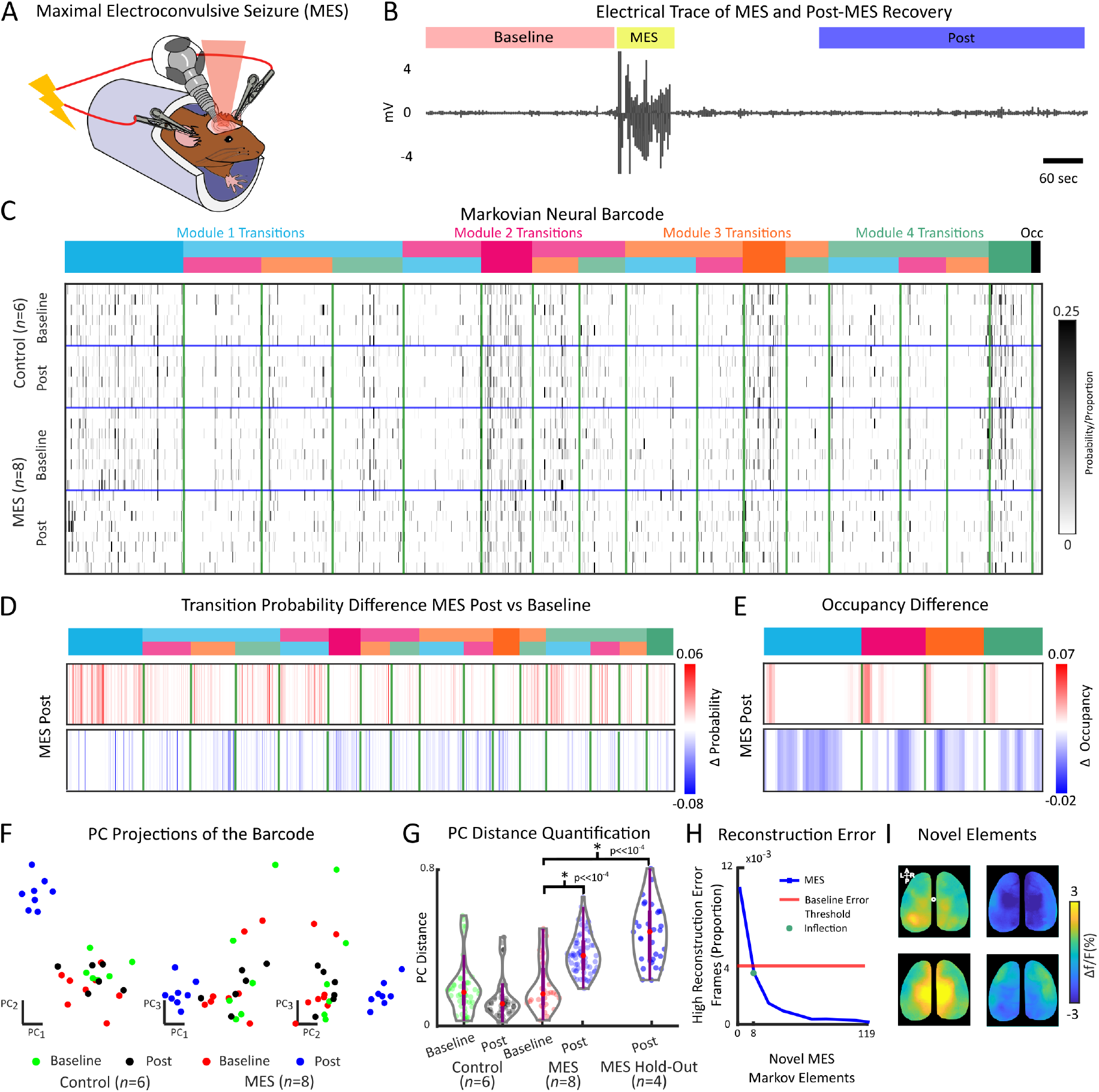
The Markovian neural barcode reveals dynamical differences and novel activity motifs after extreme excitation. **(A)** A schematic representation of the integration of the Maximal Electroconvulsive Seizure (MES) model of seizure and mesoscale cortical imaging in the awake mouse. **(B)** A voltage trace representing neural activity preceding the induction of a seizure with MES (base-line), during the seizure (MES), and following termination of the seizure (Post). **(C)** The Markovian neural barcode with colour legend. The darker colours denote higher probabilities/proportions. **(D)** To visualize the effects of the MES intervention on the Markovian neural barcode, we illustrate the mean difference between baseline-MES and post-MES to reveal up and downregulated Markov element transitions. **(E)** The mean difference between baseline-MES and post-MES reveals up and downregulated Markov element occupancies. **(F)** A principal component projection of Markovian neural barcode. The PC_1_ vs PC_2_ (Left), PC_1_ vs PC_3_ (Middle), and PC_2_ vs PC_3_ (Right) projections for the neural barcode in (**C**). The controls are plotted as green (baseline) and black (post) dots, while the MES are plotted as red (baseline) and black (post) dots. Note the clustering of the post-MES epochs in all three projections. **(G)** Statistical quantification of the principal component distance of the four conditions depicted in **C)** and **D)**, as well as an additional *n*=4 animals who received MES in a hold-out sample. The two Post-MES samples separate from the control conditions when tested against MES-Baseline (Wilcoxon Rank Sum Test; MES Baseline: 0.1667 ; MES Post: 0.3682, *p* = 5.4227*e* − 07; MES-Hold Out Post: 0.4959, *p* = 1.175581*e* − 07). Please see figure for inter-quartile range. **(H)** The continuous Markov chain reconstruction of the Post-MES epochs revealed a greater number of high-reconstruction error frames than would be expected in control conditions. Repeating the SBNNMF procedure on these frames identifies 119 Novel MES elements. The number of high-reconstruction error frames explained with each additional novel Markov element is illustrated, and the point of inflection in reconstruction improvement highlighted with a green dot and on the x-axis. **(I)** Markov element exemplars generated from the SBNNMF of high-reconstruction errors illustrating, revealing novel MES Markov elements that are not present in normative mesoscale cortical dynamics.

We therefore imaged head fixed mice (n=8) as they received MES while we sampled mesoscale cortical activity over a 5-minute baseline and 10 minute MES+Post-MES. We then constructed the Markovian neural barcode for the baseline and post-MES epochs (**Figure 5C**). To account for long exposure to LED excitation inducing photobleaching or other effects, we acquired a control recording of the same length three days prior (Control). To visualize and facilitate interpretation, we plot the difference between the group means to reveal up and downregulated Markov element transitions and occupancies (**Figure 5D&E**). This revealed an upregulation of transitions, both inter- and intra-modular, involving Module 1. Using PC projections, we see clustering of the post-MES state from other conditions (**Figure 5F**) as confirmed by PC-distance distributions (**Figure 5G**). This was also true for a hold-out validation sample of n=4 mice (**Figure 5G, SFigure 12A-C**).

As previously highlighted, a set of Markov Elements derived from a discovery sample can be applied to similar conditions with no excess in high reconstruction error frames (**SFigure 2E**). We sought to determine whether interventions, such as MES, result in novel or emergent Markov Elements as a consequence of directly manipulating the biological constraints governing transitions between Elements. In the Post-MES condition, we observed an excess of high reconstruction error frames (relative to the baseline-MES condition **Figure 5H**). When SBNNMF was applied to these high reconstruction error frames, it revealed an additional 119 Markov Elements. The point of inflection in reconstruction error frames occurred at 8 novel Markov Elements (**Figure 5H**), which was also true for the holdout sample (**SFigure 12D**). Exemplars for these novel Markov Elements emerging only after MES are illustrated in **Figure 5I**.

Evidently, extreme excitation is a powerful intervention that introduces persistent flexibility into neural dynamics and their constraints. This illustrates the power of the Markovian neural barcode as a means of discovering novel dynamics in models of pathology.

## Discussion

The spatiotemporal features of spontaneous cortical dynamics have been hypothesized to play several roles in the brain, ranging from circuit development and maintenance, to adaptive action selection^29^. However, for techniques such as mesoscale imaging and other high-dimensional methods of sampling neural activity, this signal is notoriously difficult to analyze^40^.

We propose the Markovian neural barcode as a compact lower-dimensional representation of dynamics and their constraints to resolve this impasse. This involves a novel application of discretizing cortical activity and modelling with a continuous Markov processes describing the constraints governing the brain’s dynamics as a descriptive neural barcode. This approach reveals a generalized structure across mice and acts as a formal model of brain dynamics. The SBNNMF method identifies Markovian Elements capable of discretizing mesoscale dynamics, and the associated transition probability matrices show emergent communities that serve as pseudo-absorbing states with biological significance. The Markovian neural barcode is sensitive to pathological states, as exemplified by post-ictal brain dynamics. A novel insight afforded by the approach is the identification of new and emergent cortical activity motifs that are not present under naïve conditions. Together, the method provides new avenues to determine the function of cortical motifs and understand flexibility within the constraints governing cortical dynamics.

Other unsupervised methods of identifying sequences in mesoscale imaging data (e.g. seqNMF), struggle with the dimensionality and raw file size of mesoscale imaging data. To circumvent this ‘big data’ problem, the literature to date examining activity motifs have often used experimentally generated templates to identify matching patterns of activity in recordings, as a kind of spontaneously occurring “replay”^19,20,37,38^. With some exceptions^38^, these templates have largely been sensory in nature and unsupervised methods of identifying motifs have also principally identified motifs dominated by sensory-like events^34^. Consequently, the diversity and complexity of cortical dynamics has not been fully explained by existing approaches. Parcellating spontaneous high-dimensional mesoscale cortical dynamics with Markov Elements, provides a representative set of Markov Elements to describe cortical dynamics that can be generalized to new animals and conditions. Moreover, the scalability of the method can be extended to other methods of sampling neural activity, such as functional magnetic resonance imaging, where the data structures can be several orders of magnitude larger than mesoscale cortical imaging.

The biological constraints governing cortical dynamics likely reflect a complex interplay between anatomy, neurotransmission, experience and internal state. While spontaneous fluctuations and cortical motifs have been considered internally generated in conditions of anesthesia^19,31,39^, more recent data from the awake animal indicates that behavior creates distributed activity patterns across the neocortex^23,55^. Thus, activity dynamics represent the organization of thalamocortico-signalling^62^ and neuromodulatory systems^12^ or, alternatively, it may be inseparable from the organization of animal behavior and its sequencing^46,47^. Indeed, future applications of mesoscale imaging using the Markovian neural barcode may integrate freely moving behavior with headmounted mesoscopes^63^.

Succinctly, the Markovian neural barcode is a novel method leveraging the transitions between motifs of activity to describe the constraints governing mesoscale cortical dynamics. It provides a potential solution to high-dimensional neural data generated by modern neuroscience methods, across a wide breadth of questions, to provide a novel lens to study normative and pathological brain function.

## Methods

### Animals

Primary analyses involve male and female adult C57BL/6J-Tg(Thy1-jRGECO1a)GP8.58Dkim/J^42^ (JAX #032010) mice (8-16 weeks old), constitutively expressing the red-shifted calcium indicator jRGECO1a under the Thy-1 promoter were used.

Mice were group housed on a 12:12 light cycle with ad libitum access to food and water. All Thy1-jRGECO1a and Thy1-GCaMP6F procedures were approved by the University of Calgary Animal Care Committee (ACC17-0225/ACC21-0208). All experiments were conducted in accordance with standards set by the Canadian Council on Animal Care (CCAC).

### Surgeries

Chronic skull intact window surgeries were performed as previously described^64^. Mice were isoflurane anesthetized (4% induction, 1.5-2.5% maintenance, 0.5 L/min oxygen) and buprenorphine (0.03 mg/kg) was administered subcutaneously for analgesia. Bupivacaine (Sterimax, intradermal, 0.05 mL) was administered locally at the excision site. Body temperature was maintained at 37°C with a feedback thermistor, and eyes were protected with lubricant (Opticare, CLC Medica). Following disinfection with 3x alternating chlorhexidine (2%) and alcohol (70%), the skull was exposed with a skin excision from 3 mm anterior to bregma to 2 mm posterior to lambda, and bilaterally to the temporalis muscles. A metal screw was fix to the skull with cyanoacrylate prior to embedding in transparent dental cement (C&B-Metabond, Parkell). A flat 9 × 9 mm glass coverslip (tapered by 2 mm anteriorly) was fixed to the skull with transparent dental cement, taking care to avoid the formation of air pockets. Mice recovered for 7 days prior to further interventions, allowing for full cement hardening and wound healing.

For MES experiments, electrodes constructed from Teflon-coated stainless-steel wire (178µm diameter, A-M Systems) were connected to gold plated male amphenol pins and implanted under isoflurane anesthesia as above. The electrode was implanted 750µm into posterior parietal cortex lateral to the chronic window through temporalis muscle. The implant was anchored to the chronic window using dental cement and a ground electrode, and animals were allowed to recover for a minimum of 5 days.

### Imaging Protocol

Mice were habituated to handling, head fixation with the embedded screw and the imaging apparatus including the excitation LED over 5 days.

Recordings were of different lengths for different experiments; however all acquisitions were at 50Hz temporal resolution with the exception of those intended to determine the effect of sampling frequency on the Markovian neural barcode. For experiments imaging under quiet wakefulness and for pharmacological experiments, spontaneous cortical activity was recorded for 22,500 frames (7.5 minutes). For maximal electrical stimulation (MES) seizure induction experiments neocortical calcium activity was sampled at 50Hz for 45,000 frames, corresponding to a 5-minute recording of cortical activity prior to MES, followed by 10 minutes of cortical activity after the seizure. Visual stimulation experiments involved sampling 40,000 frames of mesoscale cortical calcium dynamics.

### Thy1-jRGECO1a imaging

Cortical calcium activity during quiet wakefulness was sampled using a macroscope (Nikon 55mm lens, f/2.8 aperture) and a Quantalux 2.1 MP Monochrome sCMOS Camera (Thorlabs). We acquired 16-bit images with 19.7ms temporal resolution (50Hz) and 256×256 pixel resolution (26.5 pixel/mm). Using a 567nm excitation LED in conjunction with a 540/80 filter (Semrock) or a 470nm excitation LED attached to an articulating arm, we illuminated a wide expanse of dorsal neocortex (10-15 mW/cm^2^). Emission fluorescence was filtered with a 629/56 bandpass filter (Semrock). Focus was set ∼500µm below the cortical surface to minimize signal distortion from large blood vessels. Illumination and frame capture was controlled using commercial software (Labeo Technologies, Inc). or 15 minutes.

### Image Processing

Image stacks were analyzed using custom-written MATLAB code (Mathworks, MA). Pixel responses were expressed as a change in fluorescence relative to pixel mean fluorescence over the duration of the recording (ΔF/F0). For MES experiments, this was expressed relative to the pixel mean fluorescence over the five minutes preceding MES. The individual time-varying pixel signals were bandpass filtered (0.1-15 Hz) for analysis. MES experiments and associated controls were bandpass filtered 0.1-20Hz. Voltage sensitive dye and iGluSnFR experiments were bandpass filtered to 0.5-6Hz. Each image stack was aligned to the Allen Institute for Brain Science Mouse Brain Atlas common coordinate framework (CCF) using rigid transformation to anatomical landmarks. To restrict analysis to cortical activity during periods of quiet wakefulness, movement frames were identified as a deviation of the mean square error of spatial high-pass images from the mean image and removed prior to analyses. This corresponded to 20±5% of the total number of frames and was the only reason for excluding frames from recordings. A common mask was applied to all recordings to remove pixels peripheral to the chronic window and neocortex, as well as mask out the sagittal sinus and major tributaries.

To validate the accuracy of motion artifact measurement, a subset of mice (*n*=15) were recorded with an infrared behavior camera with mouse facial features and forelimbs in view, illuminated with a single infrared LED (Raspberry Pi night vision module) simultaneous with cortical fluorescence imaging. Motion artifacts were extracted from the full frame of the behavioral images as the mean square error of each frame relative to a 30 second moving mean image. Motion periods were identified as mean square error values >5 median absolute deviations from the median to compare with motion artifacts extracted from the fluorescence image stack. The median concordance between behavioral camera identified movement and fluorescence imaging identification of movement was 90.23% (**SFigure 17**).

### Haemodynamic Correction

A subset of mice (*n*=22) were recorded with jRGECO1a fluorescence frames interleaved with green (525nm LED) and red (625nm LED) reflectance frames for hemodynamic analysis. Red reflectance images were collected on one camera (Quantalux 2.1 MP Monochrome, 514/44 emission filter), while green reflectance and jRGeco1a fluorescence were collected on second camera (Quantalux 2.1 MP Monochrome, 618/50 emission filter), with images split with a 552nm dichroic filter. The effective frame rate for each channel was 15Hz. Frames from each camera were aligned with a rigid transform generated from a prior calibration image set. Hemodynamic correction was implemented as a pixel-wise linear regression of the df/F fluorescence signal against the dR/R reflectance signals based on established methods ^50^.

### Inter- and Intra-individual Variability in Cortical Dynamics

A subset of naïve animals were recorded repeatedly to estimate the inter- and intra-individual variability in spontaneous cortical calcium dynamics. To achieve this, animals were repeatedly imaged both within and across days and animals were returned to their homecage between recordings. Specifically, spontaneous cortical activity was sampled at baseline, after one hour, after two hours, and after 24 hours. A separate subset of naïve animals were recorded once weekly for 8 weeks.

### Maximal Electroconvulsive Shock (MES)

A suprathreshold maximal electroconvulsive shock (MES) stimulus was used to elicit a controlled generalized tonic clonic convulsion. This was delivered by a GSC 700 shock generator (model E1100DA, Grason-Stadler) through ear clips to, consisting of a 0.2 s biphasic 60 Hz sine wave pulse. As a control for MES effects on neocortical dynamics, spontaneous cortical calcium activity was sampled without any intervention two days prior to MES.

### Visual Stimulation

To determine the impact of sensory information on the structure of cortical dynamics during quiet wakefulness, we sampled neocortical activity as headfixed animals received monocular visual stimulation in the form of a full field flash. To minimize contamination of the imaging field we utilized a 450nm LED (M450LP1, Thorlabs) directed to the mouse’s right eye. We sampled cortical calcium activity animals received 200 5ms full field flashes with a 3s inter-stimulus interval and a jitter of one second. Individual and averaged visual responses were baseline corrected to the mean ΔF/F value in a 10*10 pixel ROI within primary visual cortex for the 10 frames (200ms) preceding stimulus delivery. Median split of the secondary visual response amplitude was preformed as a function of the maximal ΔF/F value of the evoked response trace within 200-400ms after stimulus delivery.

### Continuous Time Markov Chains

A Continuous Time Markov Chain (CTMC) is an extension of classical Markov chains where a stochastic system evolves continuously in time with *N* states that a system can occupy, *s*_1_, *s*_2_, … *s*_*N*_ at any moment in time *t*. The system is parameterized by two quantities, the *N×N* transition probability matrix *TPM* and a vector of *N×*1 average holding times, *μ*. Suppose at time *t* the system occupies state *i*. Then, the time that the system spends in state *i* (the holding time or occupancy time) is given by an exponential random variable in (1):

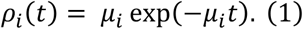

When the system exits state *i*, the system can transition to state *j* with a probability given by the *i, j*th element of the transition probability matrix, *TPM*, in (2):

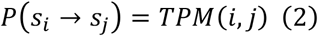

Thus, a standard CTMC can be interpreted as a set of ordered pairs (*s*_*h*(*i*)_, *t*_*i*_) where the system occupies state *h* (*i*) to *t*_*i*_ units of time, for *i* = 1,2, … *n* where *h* (*i*) : *Z*^+^ → {1,2, … *N*} is a function that denotes the *i*th state occupied. Below, we describe how the continuous time mesoscale cortical imaging recordings can be described as a CTMC with the construction of Markov Elements, that define the discrete states *s*_1_, *s*_2_, … *s*_*N*_ and the construction of the Markovian Neural Barcode, a vector used for analyzing the dynamics of a system constructed from *TPM* and *μ*.

### Supplemental Methods: Overall Sequence of Events for Constructing a Markovian Neural Barcode Using a Continuous Time Markov Chain Model

The description of creating a Markovian neural barcode to describe cortical mesoscale imaging as a continuous time Markov chain is realized by a multi-step process:

**Step 1**: Subsample a small number of frames from a selection of mesoscale recordings to create a “discovery sample”. The randomly selected frames in the discovery sample will be analyzed to determine discrete states mesoscale cortical imaging may occupy stochastically.

**Step 2:** Simultaneously apply dimension reduction and clustering to the frames in the discovery sample with semi-binary nonnegative matrix factorization (SBNMF). SBNMF decomposes the discovery sample into a set of representative features that will define Markov Elements, the states the mesoscale cortical imaging data can occupy.

**Step 3**: For each new recording, assign each frame to a Markov Element by determining the nearest Markov Element as measured by a distance metric. This time series can be analyzed as a continuous time Markov chain.

**Step 4:** Using the time series of Markov Element to frame assignments, compute the transition probability matrices (i.e. Element to Element transitions) and occupancy distributions (i.e. Element dwell time in number of frames) for the analyzed recordings.

**Step 5:** Construct and analyze the neural barcode with the transition probability matrices and occupancy distributions as a large, immediately visualizable vector. This vector can be analyzed across conditions and across mice to determine the similarities or differences in the underlying mesoscale cortical imaging recordings.

**Step 6:** Identify mesoscale imaging frames with a high error reconstruction relative to their assigned Markov Element to identify new and emergent activity motifs.

### Step 1: Creation of the Discovery Sample

For clarity, the terminology used in this section is defined in **Table 1**.

**Table 1:** Table of notations used in creating the final set of Markov Elements (the ***Discovery Markov Elements***) and optional the cross-validation controls for the Markov Elements.

| Notation | Definition |
| --- | --- |
| **Discovery** |  |
| Discovery Recordings | A subset of $n = 80$ recordings from an original $n = 160$ recordings ($n = 40$ mice imaged $n = 4$ times as presented in **Figure 3**). |
| Discovery Sample | A random sampling of 40 frames from each Discovery Recording for a total of 3200 frames. Semi-binary Non-Negative Matrix Factorization (SBNMF) is applied to the discovery sample to generate the discovery Markov Elements. |
| Discovery Markov Elements | The top 100 ranked Markov Elements by how frequently they were assigned to the Discovery Sample. |
| **Validation** |  |
| Validation Recordings | The remaining $n = 80$ recordings from an original $n = 160$ recordings ($n = 40$ mice imaged $n = 4$ times as presented in **Figure 3**). |
| Validation Sample | A random sampling of 40 frames from each Validation Recording for a total of 3200 frames. SBNMF is applied to the validation sample to generate the validation Markov Elements. |
| Validation Markov Elements | The top 100 ranked Markov Elements generated by using SBNMF on the validation sample. |
| **Random Sampling From Discovery Recordings** |  |
| Random (Discovery) Validation Markov Elements 1 | The top 100 ranked Markov Elements generated by using SBNMF on a second random sample of 3200 frames from the discovery sample. |
| Random (Discovery) Validation Markov Elements 2 | The top 100 ranked Markov Elements generated by using SBNMF 6400 frames from the discovery sample. |

To construct the set of Markov Elements, 160 recordings from *n*=40 mice (presented in **Figure 3**) was split into two randomly assigned sets of recordings, the ***Discovery Recordings*** and the ***Validation Recordings***. The Discovery recordings were used for constructing the set of Markov Elements while the validation recordings was used for cross-validating the Markov Elements. From the discovery recordings, we randomly sampled 40 frames from each recording to constitute a ***Discovery Sample*** of *n* = 3200 frames. The mesoscale imaging frames acquired for this work had a resolution of 256^*^256 pixels, and therefore this formed a 256^2^×3200 frame data matrix, that we denote *V*. Semi-Binary Non-Negative Matrix Factorization Algorithm (SBNMF) was applied to the matrix *V* (the Discovery Sample) with SBNMF to identify an initial discovery set of 150 Markov Elements. We found that 95% of the Discovery Sample were assigned to the top 100 ranked Markov Elements generated from SBNMF (**SFigure 2A**). These 100 ***Discovery Markov Elements*** were used to construct the Markovian neural barcode.

### Step 1.1: For the purposes of validating Markov Elements, additional optional steps may be taken

As a cross-validation control, we used the ***Discovery Markov Elements*** to compute the reconstruction error in the ***Validation Recordings*** for an increasingly larger number of Markov Elements, [1, 2, 3, …, 8, 9, 10, 20, 30, …, 90, 100]. This is presented in **SFigure 3**.

An identical procedure can be applied to the independent set of 80 recordings constituting the Validation Recordings, to identify the ***Validation Markov Elements*** for cross comparison. We also compared the structural similarity of the ***Discovery Markov Elements*** to the ***Validation Markov Elements*** by a Spearman rank correlation. For the ***Validation Sample***, each Markov Element was matched to a corresponding Markov Element in the ***Discovery Sample***, denoted *m*_;_. The correlation was then computed between the matched Markov Elements The significance of the correlation was calculated using MATLAB’s built in *corrcoef* function (MATLAB R2021a).

As a random permutation control, we returned to the Discovery Recordings to repeat this process with a second random sampling of 40 frames per recording to create a new 256^2^×3200 frame data matrix. The top 100 ranked Markov Elements from the subsequent application of SBNMF to this new randomly selected data set are the ***Random (Discovery) Validation Set 1***. To determine whether a larger sample altered the composition of the identified Markov Elements, this process was again repeated with a third random sampling of 80 frames from each recording in the Discovery Recordings to constitute a sample of 6,400 frames. The top 100 ranked Markov Elements from the application of SBNMF to this dataset are ***Random (Discovery) Validation Set 2***.

### Step 2: Markov Elements from Semi-Binary Non-Negative Matrix Factorization (SBNMF)

#### Semi-Binary Non-Negative Matrix Factorization (SBNMF) Algorithm

Below we will outline the Semi-Binary Non-Negative Matrix Factorization (SBNMF) Algorithm used throughout the work to discover Markov Elements. The algorithm is an adaptation of the SBNMF algorithm described by Miyasawa, Fujimoto, & Hayashi^43^. The Pseudocode for the algorithm is described in **Table 2**, while all of the parameters in the pseudocode are described in **Table 3**. Briefly, the algorithm decomposes an *m ×n*_*t*_ (*m* = *Frame dimension x* * *Frame dimension y, n*_*t*_ is the number of data frames in the sample) matrix denoted *V* as the product of two matrices as in (3)

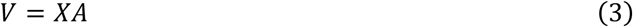

where the matrix *X* is an unconstrained (i.e. can have negative or positive elements) matrix whose columns are centroids that are to be determined, and the matrix *A* is a binary matrix

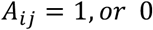

whose columns act as indicators to the centroid clusters. The matrix *X* is *Frame dimension x* * *Frame dimension y×p* while *A* is *p×n*_*t*_. In this way, for *a*_*ij*_ ∈ *A*, *a*_*ij*_ = 1 if column *X*_*j*_ ∈ *X* belongs to the *i*^*th*^ cluster, and 0 otherwise. In the algorithm, the initial number of centroids is given by the parameter *p*.

**Table 2:**
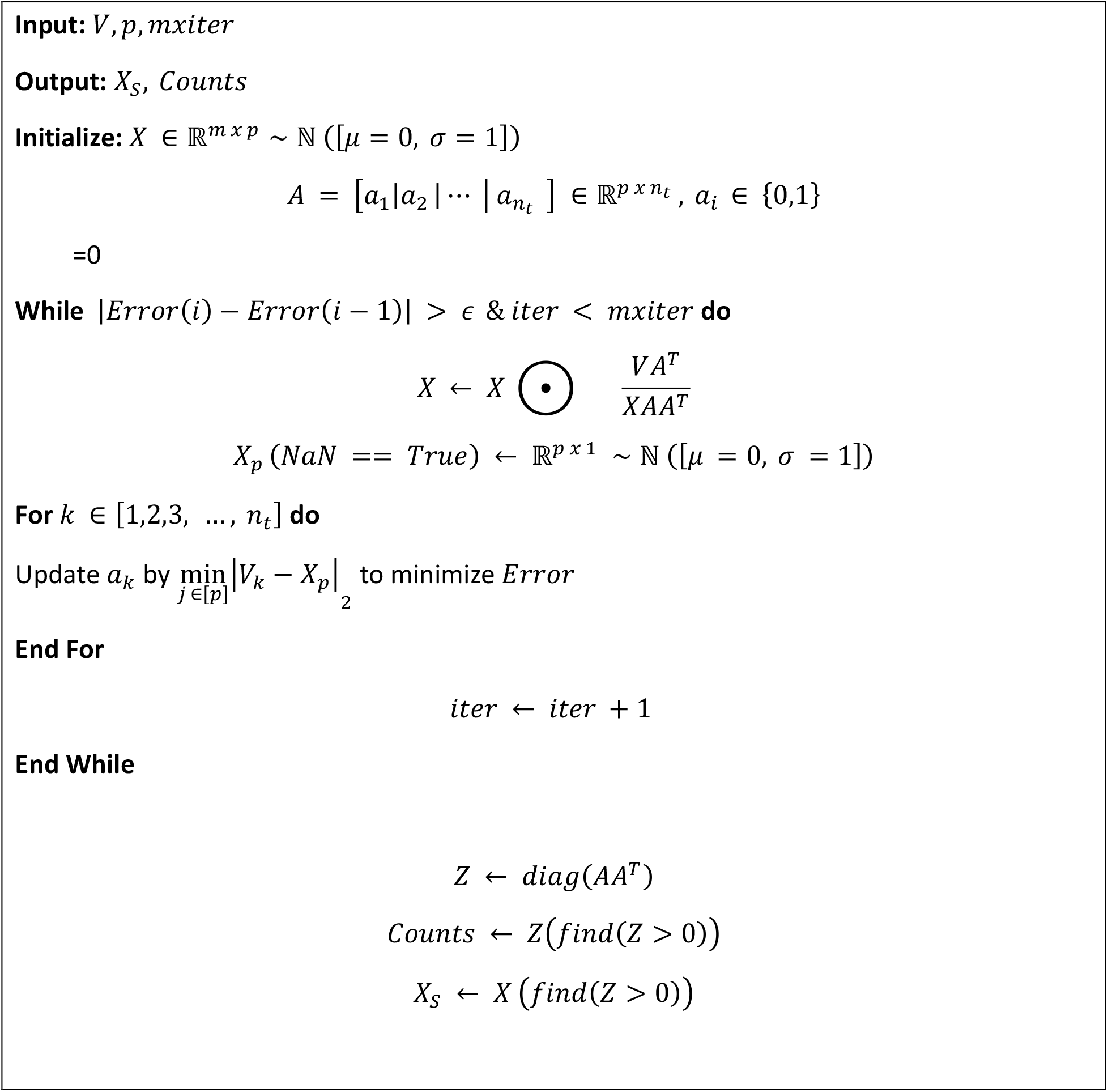
The pseudocode for the SBNMF algorithm.

**Table 3:** The parameters used their corresponding description for SBNMF to discover Markov Elements from mesoscale cortical recordings from the raw data matrix *V*.

| Variable | Description and value/range of values used in SBNMF |
| --- | --- |
| $X$ | Unconstrained $Frame\ dimension\ x * Frame\ dimension\ y \times p$ matrix whose columns are Markov Element centroids to be determined. |
| $V$ | Input data matrix that is $Frame\ dimension\ x * Frame\ dimension\ y \times n_t$ whose columns are sample frames from which Markov Elements will be derived |
| $n_t$ | Number of sample frames in $V$ |
| $p$ | Initial number of proposed centroids. |
| $\mu$ | Mean of randomly initialized centroid data |
| $\sigma$ | Standard deviation of randomly initialized centroid data |
| $A$ | Indicator $p \times n_t$ matrix of binary values. |
| $Error(i)$ | Error metric whose subsequent values are used as a convergence condition for SBNMF |
| $\epsilon$ | Error tolerance parameter used as a convergence condition for the SBNMF algorithm |
| $mxiter$ | The maximum number of iterations in the optimization algorithm used to reconstruct $V$ from $X$ and $A$ |
| $counts$ | The number of sample frames assigned to each Markov Element centroid (column of $X$). |

The SBNMF algorithm takes as input, the data sample matrix *V*, the initial number of centroids (Markov Elements) *p*, the maximum number of iterations allowed for convergence *mxiter*, and an error tolerance convergence parameter *ϵ*. The SBNMF algorithm returns as its output the sorted Markov Elements *X*_*s*_, and the associated frequencies, *Counts*, indicating how many of the initial Discovery Sample frames have been assigned to each centroid. The centroid matrix *X* is initialized from randomly generated data from a normal distribution with mean zero and standard deviation 1 for each element. The indicator matrix *A* is initialized by assigning each column of *A* to a randomly chosen standard basis vector in ℝ^*p×*1^, and the iteration count parameter *iter* is also set to zero. SBNMF iteratively seeks to minimize the difference in the reconstruction error in (4),

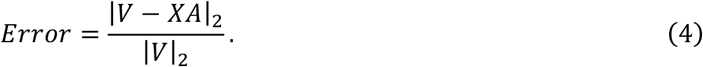

The error is minimized by updating the basis matrix *X* based on the auxiliary function technique^65^ 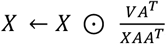, where ⊙ is the elementwise matrix multiplication (Hadamard Product), with matrix division also taken elementwise. At each update step of *X*, columns *X*_*p*_ ∈ *X* that were not assigned to Discovery Sample frames in the previous step are reinitialized so not to cause division by zero, and subsequent *NaN* entries in *X*. After each update of *X*, the indicator matrix *A* is updated in a columnwise fashion 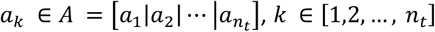, where some *j* is chosen for *a*_*jk*_ = 1, and 0 otherwise, such that 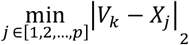 that is, *j*, is the index of the centroid that minimizes the distance to sample frame *k*. Intuitively, at each iteration of SBNMF, each column of *X*, say *X*_*j*_ is updated as the algebraic mean over the columns of *V* that have been assigned to *X*_*j*_ in the previous update of *A*. When convergence is reached, the *Counts* vector is found as the non-zero entries of the diagonal matrix product *AA*^*T*^, these non-zero entries are used to return the corresponding centroids, *X*_*S*_. The *Counts* vector in the algorithm is subsequently sorted in descending order to indicate the top ranked centroids and this ranking is applied to *X*_*S*_. Unless otherwise, stated the error tolerance parameter is taken to be *ϵ* = 0.01.

### Step 3: Markov Element Assignment

#### Markov Model Construction

The set of Discovery Markov Elements is denoted as the set B. To construct a Markov chain model, M, for a given recording, R, with *n*_*t*_ frames, each data frame, R_i_, *i* ∈ [1,2, …, *n*_*t*_], was paired with a corresponding global Markov Element, B_*k*_, *k* ∈ 1,2, …, 100, by finding the index *k* that solves the minimization in (5):

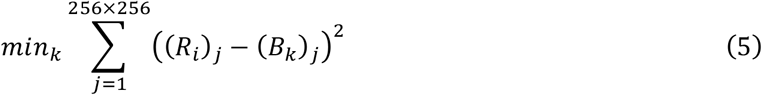

The index, 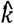, that minimizes the sum of squares difference above in (5), is the assigned Markov element, 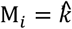, for frame R_*i*_ of R.

#### Cost Minimization, Optimal Lambda, and Flicker Optimization

To remove spurious boundary ‘flicker’ in our Markov Element reconstruction, wherein mesoscale cortical activity is nearly equidistant between two Discovery Markov Element centroids introducing ‘flicker’ transitions, we seek to optimize the cost metric given in (6):

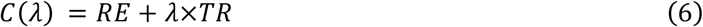

for *λ* as a trade-off between the number of transitions, *TR*, and the reconstruction error, *RE*. Here, 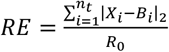 is defined as the ratio of the sum of framewise reconstruction error to the original framewise sum *R*_0_, with *X*_*i*_ the *i*^*th*^ frame in a recording and *B*_*i*_ it’s assigned Discovery Markov Element. Likewise, 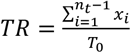 is defined as the ratio of the sum of total transitions to the number of transitions in the original assignment, *T*_0_. Here, *x*_*i*_ is defined as 1 if a transition occurs from index *i* to index *i* + 1, and 0 otherwise. For each Discovery Recording, 50,000 iterations were run, in which on each iteration an index, *i*, was randomly drawn, with corresponding Discovery Markov Element assignment *m*_*i*_. The adjacent Markov Elements assignments, [*m*_*i* − 1_, *m*_*i*_, *m*_*i* + 1_], are then checked for a flicker transition. A ‘flicker’ transition is said to occur for assignment *m*_*i*_ if (*m*_*i* − 1_ = *m*_*i*_, *m*_*i*_ ≠ *m*_*i* + 1_] or vice versa. If a ‘flicker’ transition exists, the ‘flicker’ is removed by assigning the frames in indices *i* − 1, *i, i* + 1 to the same Markov Element (assigned to *m*_*i*_) The *RE* and *TR* are recomputed after each iteration. After the removal of each ‘flicker’ transition the value of *TR* decreases and the value of *RE* increases, leaving the optimal *λ, λ*_*opt*_, as the proportion of ‘flicker’ transitions removed to minimize the cost function. The *λ*_*opt*_ was computed for our Discovery Recordings with a mean value of 0.5842 ± 0.0783 and for the Validation Recordings with a mean value of 0.5782 ± 0.0836. Thus, *λ*_*opt*_ was fixed at 0.58.

In the subsequent Markov Construction (as otherwise stated) we apply our optimization algorithm with the optimal *λ*_*opt*_ as outlined above. In this optimization algorithm, an index *i* is randomly drawn, with the corresponding Discovery Markov Element assignment *m*_*i*_. The adjacent Markov Elements assignments, [*m*_*i* − 1_, *m*_*i*_, *m*_*i* + 1_] are then checked for a ‘flicker’ transition. If a flicker transition exists, the flicker is removed by assigning the frames in indices *i* − 1, *i, i* + 1 to the same Markov Element (assigned to *m*_*i*_). This process continues until the optimal *λ*_*opt*_ is reached where the number of transitions is counterbalanced by the reconstruction error for each recording.

### Step 4: Estimation of Transition Probability Matrices/Occupancy Distributions

#### Estimation of Transition Probability Matrices and Occupancy Distributions

The Markov chain model provides several levels of information.

First, it provides an unconditional description of cortical activity states through proportional distribution of Markov Elements during a given mesoscale cortical activity recording as provided by the occupancy distribution. Here, a consistent estimator of the occupancy distribution is given from the distribution of observed frequencies in the Markov chain. The unconditional probability for the *i*^*th*^ state is given by (7):

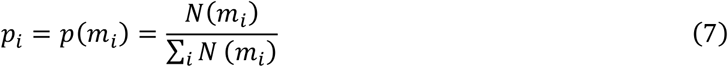

where *m*_*i*_ is the *i*^*th*^ element in the 100 element state space, and *N*(*X*) counts the occurrence, or frequency, of element *X* in the Markov chain model. In the construction of the occupancy distribution, contiguous frames that assigned to the same Markov Element, so called ‘self-transitions’, are considered.

Secondly, from this model the conditional probabilities are derived for the corresponding transition probability matrix, *P*, as found by the maximum likelihood estimates^42^ in (8):

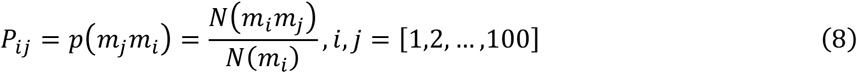

In the case where state *m*_*i*_ does not occur in the chain, it is assumed that *p*_*ij*_ = 0, *j* ∈ [1, …, 100]. In the construction of the transition probability matrix, only transitions between Markov Elements are considered. ‘Self-transitions’, or repeated Markov Element assignments within the Markov chain, are not considered in computing the transition probability matrix.

Note that for a Markov chain, the empirical estimator of a transition probability matrix, *P*, has an associated estimator of the stationary distribution *π*, where *π*_*i*_ = *p*_*i*_ is the unconditional probability from above. It can be shown that this empirical estimator, *π*, does not strictly satisfy the equation, *π* = *πP*, but for a sufficiently large number of observations in the Markov process, under certain conditions, the error is taken to be small. Here, estimators with and without self-transitions (occupancy distributions and transition matrices respectively) are considered. For the purposes of formulating the Markovian neural barcode as incorporating all of the generating/descriptive statistics of the underlying continuous time Markov chain assumption, self-transitions statistics were incorporated into the associated occupancy distributions that becomes integrated into the neural barcode.

#### Louvain Sorting Of Transition Probability Matrices

Louvain sorting of Transition Probability Matrices was done using the *Community_Louvain*.*m MATLAB R2021a* package, which is a generalization of the Louvain community detection algorithm.^66^ This function treats the averaged Transition Probability Matrix from the Discovery Recordings (**Figure 1H**) as the connection matrix for an underlying graph network. The returned community structure is one in which nonoverlapping groups of nodes, or transitions, are sorted based on maximizing the number of within-group edges, and minimizing the number of between-group edges. In this work, a uniform resolution parameter value of *γ* = 0.75 was used for all modeling to minimize singleton and doubleton modules, unless otherwise stated. The resolution parameter, *γ*, for the Louvain community detection algorithm, affects the size of the recovered modules in the Louvain algorithm. Smaller resolution values recover larger modules and therefore fewer number of modules overall, while, conversely, larger values of the resolution parameter recover modules containing fewer Markov Elements. This resolution parameter may require customization based on the activity indicator utilized, the biological source of its signal and its kinetics to minimize singleton and doubleton modules (see **SFigure 11** for realizations with singleton modules).

### Step 5: Construct and analyze the Markovian neural barcode

#### Markovian Neural Barcode Construction

The Markovian neural barcode in our analysis was systematically constructed by unwrapping the transition probability matrix (100*×*100) for each recording into a vector (1*×*100^2^) and concatenating it’s associated occupancy distribution (1*×*100). Thus for an analysis with *k* recordings, the corresponding neural barcode is of size *k×* 100^2^ 2 100 = *k×*10,100. For computational and visual convenience, recordings representing a common condition were grouped together in the corresponding neural barcode and separation between groups delineated with a blue line. For plotting clarity, columns of the neural barcode were sorted by Module as defined by the Louvain Sorting Algorithm, with a corresponding color-coded legend to facilitate interpretation of intra-module and inter-module transitions.

#### PC-Distance Matrix and Intra-Inter Group Distances

Comparative analysis of the recordings using their associated transition probability matrices and occupancy distributions, as represented in the Markovian neural barcode, was done using an *intra-inter* group analysis.

For this *intra-inter* group analysis, principal component analysis was first applied to the relevant Markovian neural barcode(s), and the first three principal components for each recording were stored in a new reduced coefficient matrix. A pairwise distance matrix, say *M*, was then calculated from the rows of the previous reduced principal component matrix using the Euclidean norm.

To quantify the separation in principal component space between a baseline group, *G*_*B*_, and condition group, *G*_*C*_, the corresponding *intra-inter* group distances can be found from *M*. If the rows of *M* that correspond to the baseline recordings are *b*_1_, *b*_2_, …, *b*_*n*_ then the *intra*-group distances are the collection of values defined by *M* (*b*_*i*_, *b*_*j*_) for *i, j* = 1,2, …, *n*, where repeats (*M* (*b*_*i*_, *b*_*j*_)= *M* (*b*_*i*_,*b*_*j*_)), and self comparisons (*M* (*b*_*i*_, *b*_*i*_)) not used in estimating the distance distributions. If *c*_1_, *c*_2_, …, *c*_*m*_ are the rows of *M* corresponding to the recordings of the condition group, then the *inter*-group distances are the collection of values defined by *M* (*b*_*i*_, *c*_*j*_)for *i* = 1,2, …, *n* and *j* = 1,2, …, *m*, where again repeats, self comparisons, and also *intra*-condition values (*M* (*c*_*j*_, *c*_*k*_), *j, k* = 1,2, …, *m*) are ignored.

The significance of *intra-inter* group distances was computed by the Wilcoxon Rank Sum test. The distance was considered at a 5% significance level, with exception where, due to multiple comparisons, the significance level was adjusted using the Bonferroni correction.

### Step 6: Calculate the reconstruction error to identify new and emergent cortical activity motifs

#### Markov Element Reconstruction Error (*L*_2_)

The reconstruction error associated with the assignment of a Markov Element as a centroid best representing a frame of mesoscale cortical activity is taken as the relative error in the matrix (vector) 2-norm. For a given recording *R*_*j*_ (or data frame) and the matrix of assigned Markov Elements *S*_*j*_ (or assigned Markov Element per data frame) the reconstruction error would be given as (9):

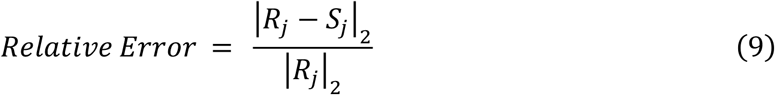

Where for the *n×m* matrix *x*, the matrix 2-norm is defined as the largest singular value of x

#### High Reconstruction Error (HRE) Frames and Condition Specific Novel Markov Elements

Under naïve conditions (i.e. no experimental manipulation other than removing mice from their homecages for headfixation and mesoscale imaging), Discovery Markov Elements describe Validation Recordings with the expected rate of high reconstruction error frames, which we arbitrarily define as >3 *σ* above the reconstruction error observed in the Discovery Sample. Therefore, the emergence of high-reconstruction error frames under experimental manipulations exceeding this threshold identifies new cortical activity motifs not found under naïve conditions, to which SBNNF can be applied to identify these motifs.

High Reconstruction Error (HRE) frames for an experimental condition were found by comparing the distributions of pooled element-wise reconstruction error data from the given condition against the pooled element-wise error from our combined discovery and validation sets, which was used as a control. These pooled element-wise distributions of reconstruction error were formed by collecting in an element-wise fashion the reconstruction error from the Markov construction process across recordings. For each Markov Element, [1…100], the pooled reconstruction error data from the experimental data and control data were both *z* scored 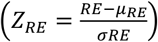 against the control data for that Markov Element. The subsequent normalized distributions were then tested using a right-tailed two-sample t-test for unequal variances to test the null-hypothesis that the reconstruction error data comes from a population with equal mean. For each significant test, the experimental condition data frames that have a normalized score greater than 3 *σ* are collected as High Reconstruction Error (HRE) frames.

The comparative analysis of animal recordings under variable experimental protocols allows for the derivation of novel, condition specific Markov Elements (nME). We use the experimental condition specific HRE frames as a sample input matrix for the SBNMF algorithm (error tolerance of 0.001, maximum number of iterations of 100, and initial number of Markov Elements = 150). The ranked nME from this application of SBNMF are then used to compute the HRE versus nME curve. The HRE versus nME curve is computed by calculating the number of subsequent HRE frames that would be found as more ranked nME from SBNMF are included in the overall Markov Element set. The number of novel Markov Elements that correspond to the inflection point in the HRE versus mME curve is then used as the initial size of the final novel condition specific set of Markov Elements. The inflection point of the HRE vs nME curve is interpreted as the optimal point to which additional nME no longer meaningfully explain condition specific motifs.

## Additional Computational Methods

### Average Histogram of Markov State Occupancy Across Time

Markov models from visual evoked response recordings (16 trials, *n* = 8 *animals in* 2 *trials each*) were aligned across stimulation trigger times (100 per trial) to a common time point *t*_*c*_ = 0, with null value padding added to time points which were removed due to motion artifact. The relative frequency of each Markov Element was then computed at each aligned time point, plus or minus 1 second around the aligned trigger time *t*_*c*_ = 0. The subsequent relative frequencies of each Markov Element were then sorted at each time point as per the previously determined Louvain sort in Figure 1.

### Application of Vessel Artifact

The vessel artifact masks were manually extracted for *n* = 3 mice. Each corresponding mask was then scaled to a maximum fluorescence value of 0.2 ΔF/F0 for mask entries and zero otherwise. The corresponding mask was then added to each frame of mesoscale cortical imaging acquired from other animals to systematically introduce a vessel artifact.

### Order Estimation of Markov Models

To estimate the order of the Markov process in the mesoscale cortical imaging recordings, the Bayesian Information Criterion^41^ (*BIC*) was computed for each recording in the repeated acquisition experiments (**Figure 3**). The equations for the log-likelihood (*LL*_*k*_)for the *k*^*th*^ order (10), the likelihood ratio statistic (_*k*_*η*_*m*_)(11), and the *BIC* for order *k* used in the order estimation (12) are shown below. For the BIC test, the objective is to find an optimal 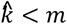 that minimizes the *BIC* criterion Here, *m* is a fixed upper bound parameter for the order of the Markov chain.

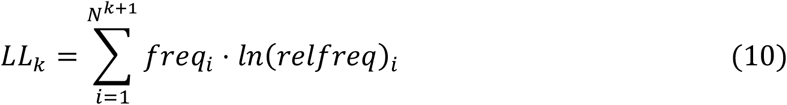

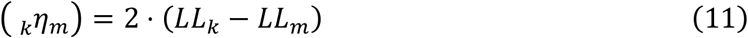

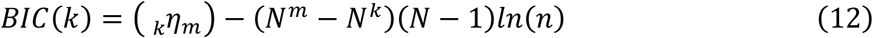

*N* = 100 is the number of Markov Elements of the state space, *n* is the number of observations in a given Markov chain, *freq*_*i*_/ *relfreq*_*i*_ is the frequency/(relative frequency) of the *i*^*th*^ element in the set of *k*^*th*^ order parameters. We note that all self-transitions are removed in the application of the BIC tests.

Furthermore, we take the convention that *n* ≫ *N*^*k*^, to consistently estimate the model parameters *freq*_*i*_ and *relfreq*_*i*_ under the assumption of a *k*^*th*^ order model. Thus, given the lengths of our recordings we take *m* = 2 and test against a zeroth and first order Markov model. Additionally, due to the size of our state space, *N* = 100, we take the convention that the number of Elements in our degrees of freedom computation, *N*^*m*^ − *N*^*k*^ is simply the number of observed Elements under the (*k*^*th*^, *m*^*th*^) order paradigm.

### Lorenz Attractor – Continuous Dynamics Model

To apply Markovian neural barcoding to continuous dynamics, we used the system of ordinary differential equations (13) known as the Lorenz system^67^ as given below.

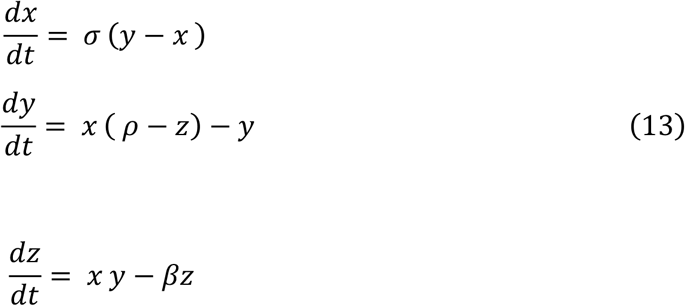

This system classically displays many chaotic attractors that subtly change depending on the specific parameter set used. We simulated the Lorenz system for two parameter sets that were nearby yet yielded slightly different dynamics and attractor shapes. For Parameter Set 1, the parameter values were set to *σ* = 10, 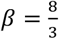, *ρ* = 28, while for Parameter Set 2, the values were set to *σ* = 10, 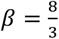, *ρ* = 32. We simulated 10 trials with each parameter set with a total simulation time of 300 seconds, with a step size of 0.001 seconds with an explicit Runge-Kutta 45 integrator. The Lorenz Discovery Sample for this system was randomly drawn from 1% of the total data points from all 20 simulations. SBNMF was then applied to the Lorenz Discovery Sample to generate an initial set of 300 Markov elements, with error tolerance of 0.001, and a maximum number of iterations set to 50..

### Continuous Time Markov Chain (CTMC) Model Parameter Estimation

The exponential transition time parameter estimation of our Continuous Time Markov Chain (CTMC) model was done using the 40 baseline condition recordings from our repeated acquisition protocol (**Figure 3**). For each baseline recording *j*, the Markov construction algorithm was applied and a Markov model, *mc*_*j*_, was constructed, along with a corresponding Markov model 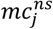 without self transitions, a separate vector of self-transition counts 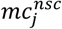. Here, if *mc*_1_ = (1,1,1,2,3,3,1,1, …), then we would have 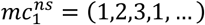, with 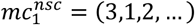. To estimate the exponential transition time for each Markov Element, 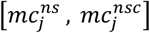 vectors were concatenated and compiled for all 40 Markov models and a distribution of counts, *C*_*k*_, from the column data of 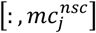, values were collected for each Markov Elements, *k* ∈ [1,2, …, 100], present in column 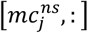. The *C*_*k*_ values for each Markov Element was then fit by the exponential model parameter function (*expfit* MATLAB R2021a) to estimate the transition time for that Markov Element. Note that the exponential transition time parameters was scaled by 50, to account for our 50 Hz sampling rate. The root *j*^*th*^ square error (RMSE) for each exponential transition time fitting for the Markov Element was taken as the RMSE between the exponential pdf (*exppdf* MATLAB R2021a) and the normalized histogram (*histogram* MATLAB R2021) *of C*_*k*_ values. To compute the “Within Recording Mean Time Between Transitions” (**SFigure 4**) the average dwell time spent in each Markov Element (average self-transition count divided by 50) was computed for each Markov Element in a single recording and then averaged across elements.

### Transition Probability Matrix Convergence and Correlation Computation

The convergence of the 100*×*100 Transition probability matrix (TPM) for a given recording was computed as the relative error in the matrix 2-norm between the TPM of the full Markov model and the TPM of abbreviated models of increasing time index. Briefly, the TPM matrix, *TPM*_*n*_ (*i, j*) is estimated using an increasing number of transitions, *n*. The matrix 2-norm *(norm(-,2)* MATLAB 2021) is used to assess the similarity between the TPM matrix with *m* transitions vs. *n* transitions with (14)

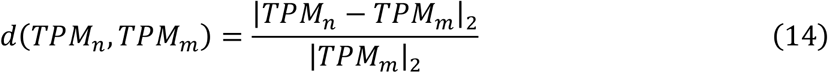

Where *m* is taken as the value that generated the full Markov model. In **SFigure 9A**, the full Markov model was taken to be 15,000 frames uniformly for each recording.

As a second set of comparison measures, the correlation between the transition probability matrices in the Discovery Sample under two different simulation conditions was computed. Under the first simulation condition, the rows of each transition probability matrix were randomly shuffled and under the second simulation condition, the transition probability matrix for each recording was computed with the Markov elements shuffled in time.

### Variance Explained

To quantify the accuracy in the lower dimensional representation of the recordings, the cumulative and individual variance explained was computed across an increasing number of principal components. To calculate these variances, we first applied the singular value decomposition (*svd*, MATLAB, R2021a) individually to the 80 recordings from our discovery sample. For each corresponding set of singular values, *λ*_1_, …, *λ*_*k*_,the individual percentage of variance explained for a specific singular value, *λ*_*j*_, is given by (15):

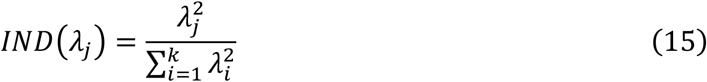

To find the cumulative variance explained for the first *j* principal components, the equation (16) was used:

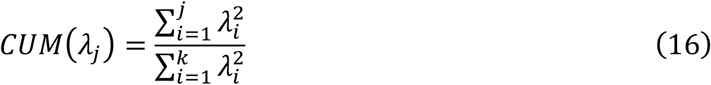

In a similar fashion, the same formulas were applied to test for the total variance explained across principal components in a lower dimensional representation of the neural barcodes.

### Difference/Standard Deviation Heat Map Barcode

Difference Heat Maps for transition probability matrices and their associated occupancy distribution were computed by first averaging individual neural barcodes across conditions and then taking the difference against a chosen baseline condition, i.e. *Condition – Baseline*. The resultant barcode of average conditional differences was then sorted by positive and negative values and plotted. Each positive and negative difference barcode was thresholded to 2 standard deviation above the mean. This process was done separately for the occupancy distribution in addition to the full Neural Barcode. The standard deviation barcode was taken as the standard deviation of conditional average barcode as defined above. The standard deviation was computed on an individual transition/occupancy basis. A 2D Gaussian smoothing filter (*imgaussfilt*, MATLAB, R2021a) with standard deviation of 1 was apply to each Heat Map before plotting.

## Acknowledgements

This study was funded through the Campus Alberta Innovates Program Chair in Neurostimulation (AM) and Discovery Grant from the Natural Sciences and Engineering Research Council of Canada (RGPIN-2020-05273). JMC received support from the Pacific Institute for the Mathematical Science. DMA received support from the Canadian Open Neuroscience Platform. AGG received support from SUDEP Aware. GCT received a grant from the Natural Sciences and Engineering Research Council of Canada (RGPIN-2019-04140). WN received support from a Hotchkiss Brain Institute Start-Up funds and a Natural Sciences and Engineering Research Council of Canada (RGPIN-2020-00334).

