## Supplemental Figures for "A Markovian neural barcode representing mesoscale cortical spatiotemporal dynamics"

### Supplemental Material

#### Videos

**Supplemental Video 1. An example of Thy1-jRGECO1a mesoscale cortical activity.**

**Supplemental Video 2. Reconstructions of mesoscale calcium imaging with increasing number of Markov Elements.**

**Supplemental Video 3. Synthetic data generated from experimental transition probability matrices.**

**Supplemental Video 4. The Lorenz system.**

**Supplemental Video 5. Individual and averaged examples of visual evoked responses.**

**Supplemental Video 6. An example of Thy1-jRGECO1a mesoscale cortical activity before and after MES.**

### Figures

##### Supplemental Figure 1. Mesoscale cortical imaging occupies a low-dimensional space.

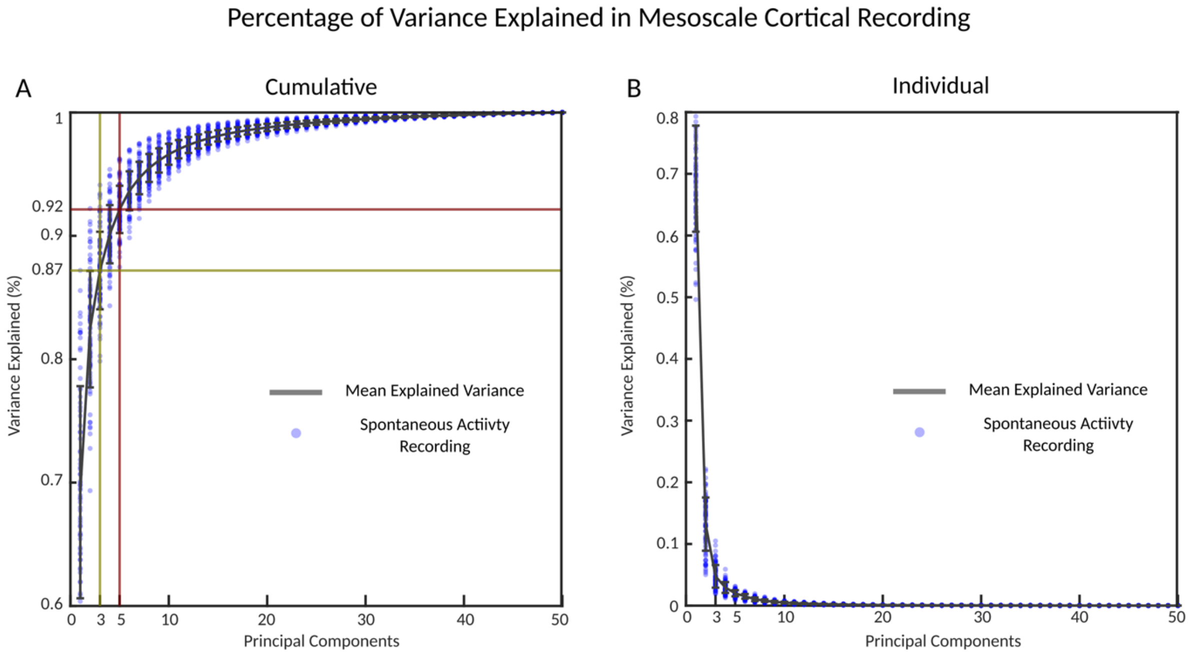


**(A)** The cumulative variance explained by principal components in a low dimensional representation of the ($n = 80$ ) randomly selected spontaneous activity recordings as given in our discovery set (**Figure 1, SF1**). Approximately 87% of the cumulative variance can be explained with the first three principal components and approximately 92% of the explained variance with the first 5 principal components.

**(B)** The individual variance explained by principal components similarly shows that the first principal components explain the majority of the explained variance.

**Supplemental Figure 2. Generation of the Markov Elements.**

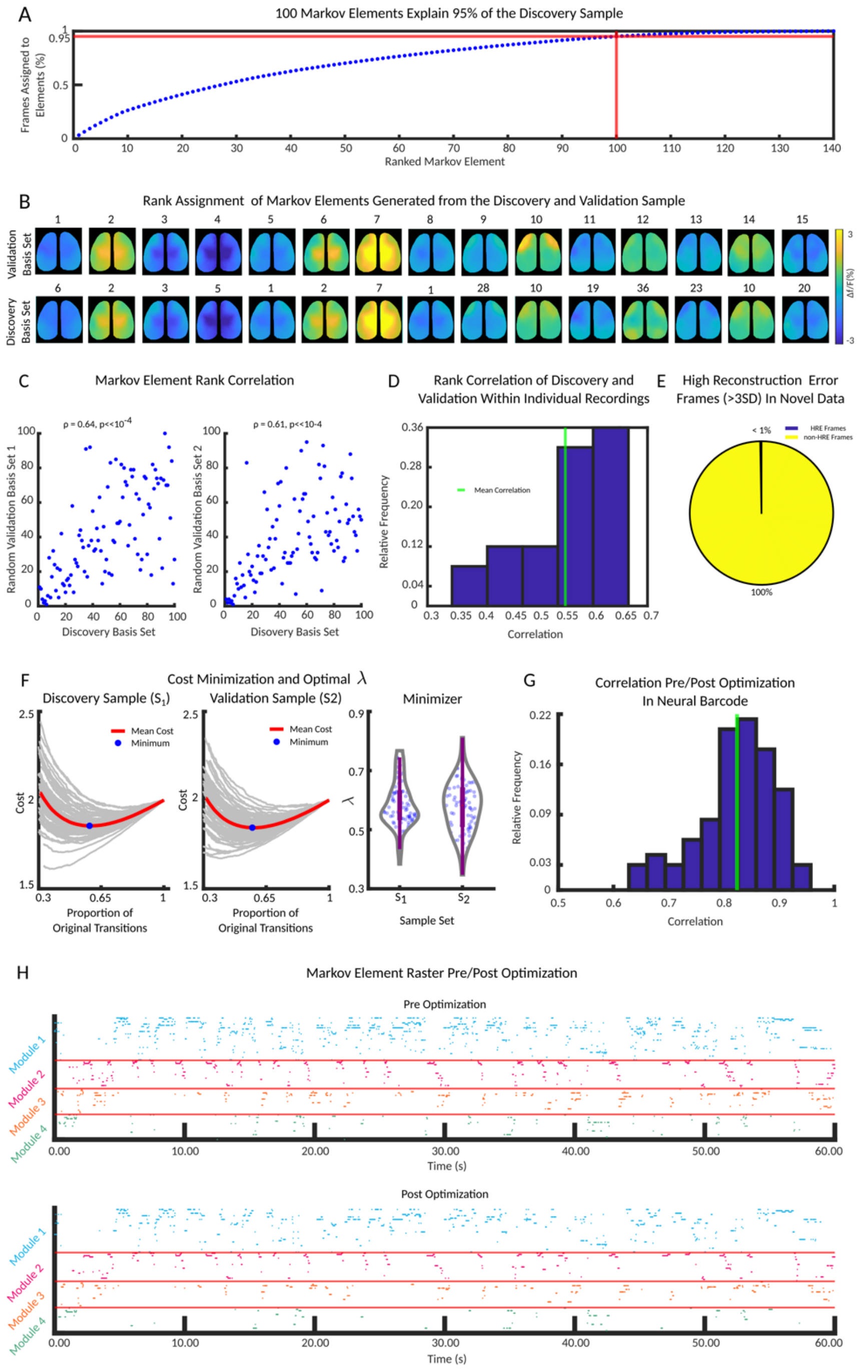

(A) The cumulative distribution of discovery sample frames ($n = 3{,}200$) assigned to the ranked Markov elements from SBNMF as a function of increasing number of Markov Elements. We find that 95% of the discovery sample can be explained with 100 top ranked Markov elements.
(B) The first 15 Markov elements of the validation set (top) with the most similar Markov elements of the discovery set (bottom), where the similarity was measured by the Euclidean distance. Note that the highest ranked Markov elements of the discovery set tended to match with the highest ranked elements of the validation set (Spearman's rank correlation $\rho = 0.64$, $p = 8.423e - 13$).
(C) The rank of the discovery basis set versus the rank of the closest matched Markov element in the first random validation basis set (left) and a second random validation basis set derived using twice the number of frames from each recording (right). The Spearman's rank correlations were $\rho = 0.64$ (left), and $\rho = 0.61$ (right). Both Spearman correlations were statistically significant ($p = 4.580e - 13$, $p = 1.36e - 11$ ).
(D) At its extreme, a basis set can be generated from a single recording from a single animal. Depicted here are the distribution of correlations of the rank frequency of elements generated from a single recording ($n = 25$ random permutations) and the discovery Markov Element basis set (mean$\pm$std correlation: $(0.5506 \pm 0.0877)$).
(E) Pie chart demonstrating the percentage of frames with high reconstruction error (HRE, blue) relative to non high-reconstruction error frames (yellow) when applying the discovery Markov Element basis set to the cross-validation set of recordings ($n = 80$ recordings; HRE/(Total Frames): 4,179 / 1,425,659).
(F) Results of the application of our cost minimization function to reduce spurious transitions (flicker) in the Markov constructions of the mesoscale recordings in the discovery sample (left) and validation set (middle). Trajectories for individual recordings (gray) are averaged (red) and the minimal value of this curve as a function of the proportion of original transitions is given in blue (left: $0.5844$, right: $0.5723$). The distribution of cost minimization values, $\lambda$, (right) is given for each sample set, the median (mean $\pm$ std) values of each sample are $0.5636$ $(0.5842 \pm 0.0783)$(discovery set $S_1$, $n = 80$) and $0.5819$ $(0.5782 \pm 0.0836)$ (validation $S_2$, $n = 80$). Please see figure for inter-quartile range. From here we choose our optimal $\lambda$, $\lambda_{opt}$, to be $\lambda_{opt} = 0.58$.
(G) The histogram of correlation coefficients between the Markovian neural barcodes for the repeated imaging protocol ($n = 160$) under regular reconstruction, and under the spurious transition minimization (flicker) protocol. The mean $\pm$ standard deviation for the correlation coefficient was $0.8301 \pm 0.069$
(H) The "raster plot" for 1 minute of mesoscale cortical imaging. The raster plot is computed by using the discrete time series of Markov element occupancies, $(t_k, M(t_k))$ where $t_k$ is the $k$th time unit, and $M(t_k)$ is an integer between 1 and 100, denoting the Markov element assigned to the frame at time $t_k$. A point is plotted at $(t_k, M(t_k))$ and colour coded by the Louvain sorted modules. The raster is shown without flicker-minimization (top) and with flicker minimization (bottom).

**Supplemental Figure 3. Increasing number of Markov Elements improves the reconstruction error.**

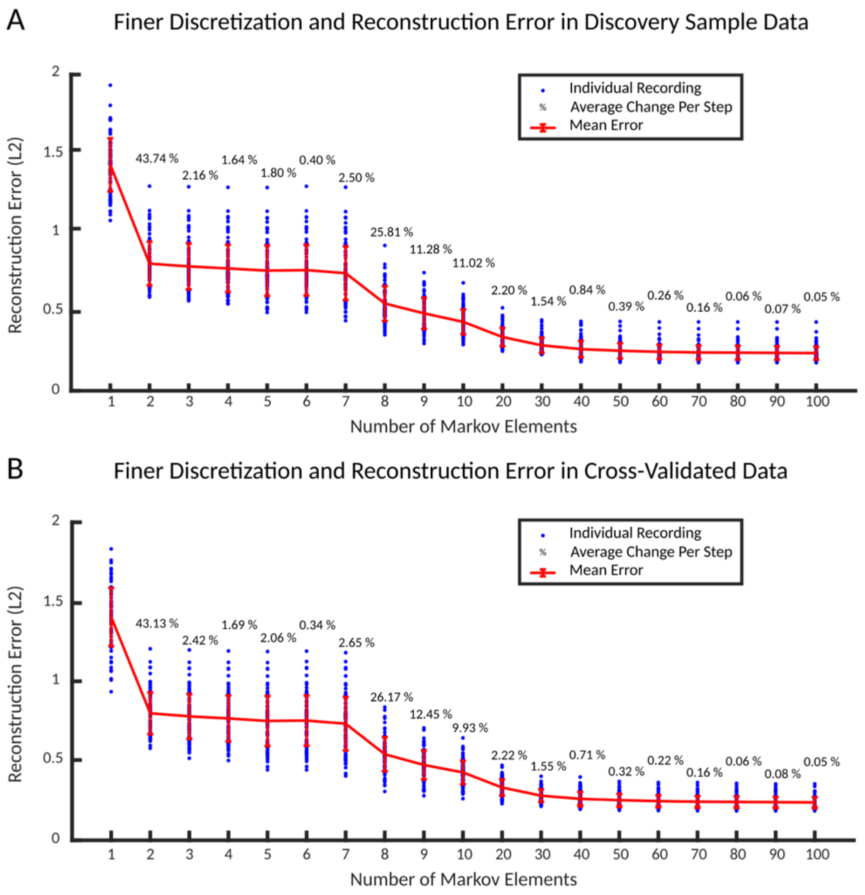


(A) The proportional change in reconstruction error for the discovery set, ($n = 80$), as more Markov Elements (from the discovery set) are used in reconstructing recordings. The reconstruction error (y-axis) is measured as the relative error in the matrix 2-norm between a recording and the Markov element reconstruction. The first 30 Markov elements contribute to the majority of the reconstruction accuracy for the discovery set.

(B) The proportional change in reconstruction error for the validation data set, ($n = 80$), as more Markov Elements (from the discovery set) are used in reconstructing recordings. The reconstruction error is measured as the matrix 2-norm between a recording and the Markov element reconstruction. Similar to the results in (A), the first 30 Markov elements contribute to the majority of the reconstruction accuracy for the validation set.

**Supplemental Figure 4. Markov Elements transition over time.**

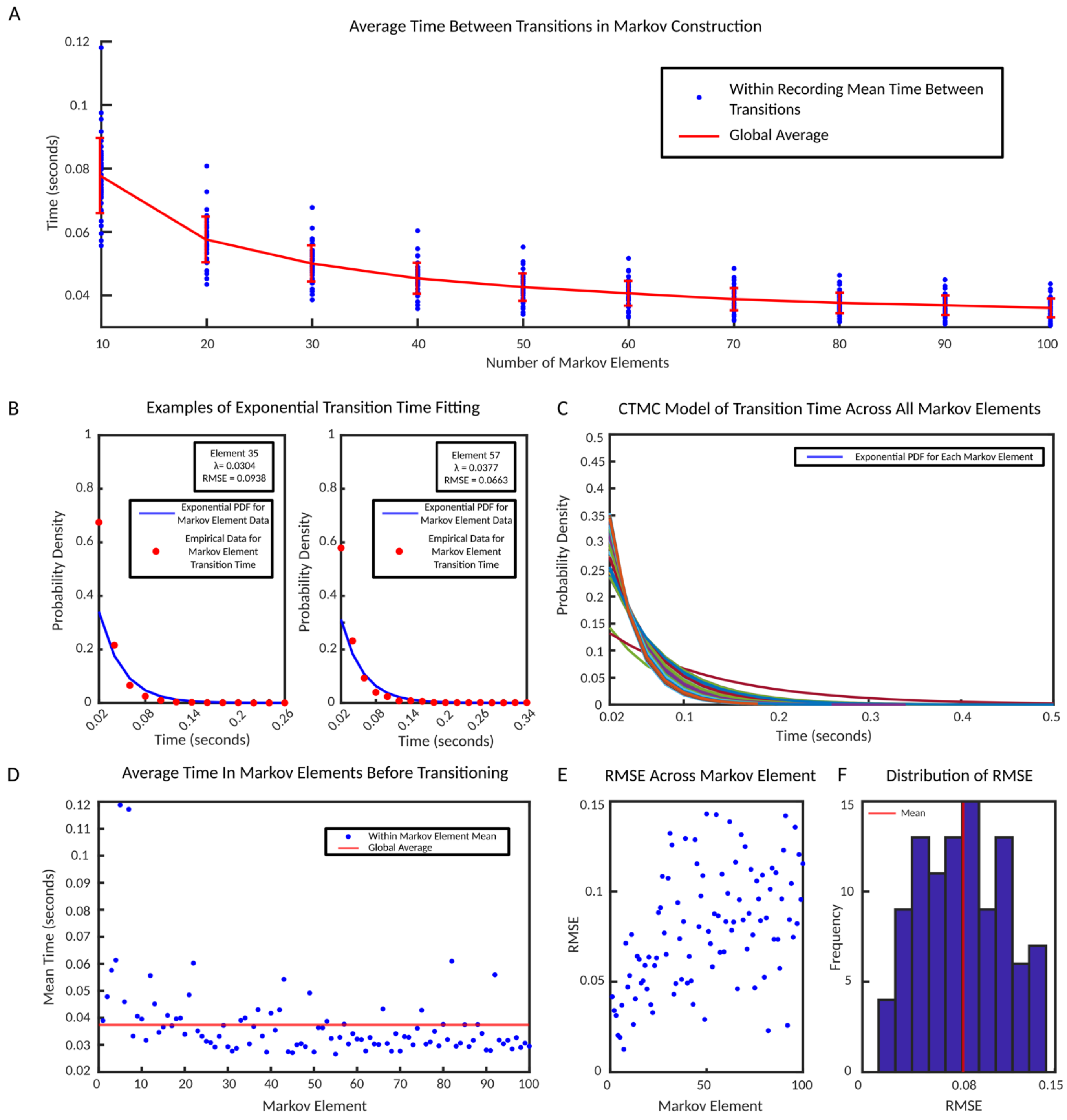

(A) The average time spent in a Markov Element by the 40 baseline recordings in the repeated imaging protocol . The average time (blue) is computed in each recording as the average dwell time across all Markov Elements for a single recording as more Markov Elements are added in the data reconstruction. Note that as additional Markov Elements are utilized in the reconstruction, the average time between transitions decreases.
(B) The empirically estimated dwell time (red points) versus the exponential fit (solid blue line) predicted by a continuous time Markov chain for a pair of Markov Elements; Element 35 (left) and Element 57 (right).
(C) The empirically fit exponential probability density function for the Markov Element dwell time across all 100 Markov Elements.
(D) The average time spent in a particular Markov Element when all 100 Markov Elements are used in the reconstruction (without flicker minimization). Note that the global average (red) is approximately $(0.0374 \pm 0.0141)$ (mean$\pm$std) seconds.
(E) The root-mean-squared error (RMSE) for each Markov Element exponential transition time fitting. The RMSE was computed as the square root of the squared error between the empirical data for Markov Element transition time and the associated exponential transition time fitting.
(F) The distribution of the RMSE pooled across Markov Elements (mean$\pm$std $(0.0785 \pm 0.0328)$).

**Supplemental Figure 5. The Markovian neural barcode also occupies a low-dimensional space.**

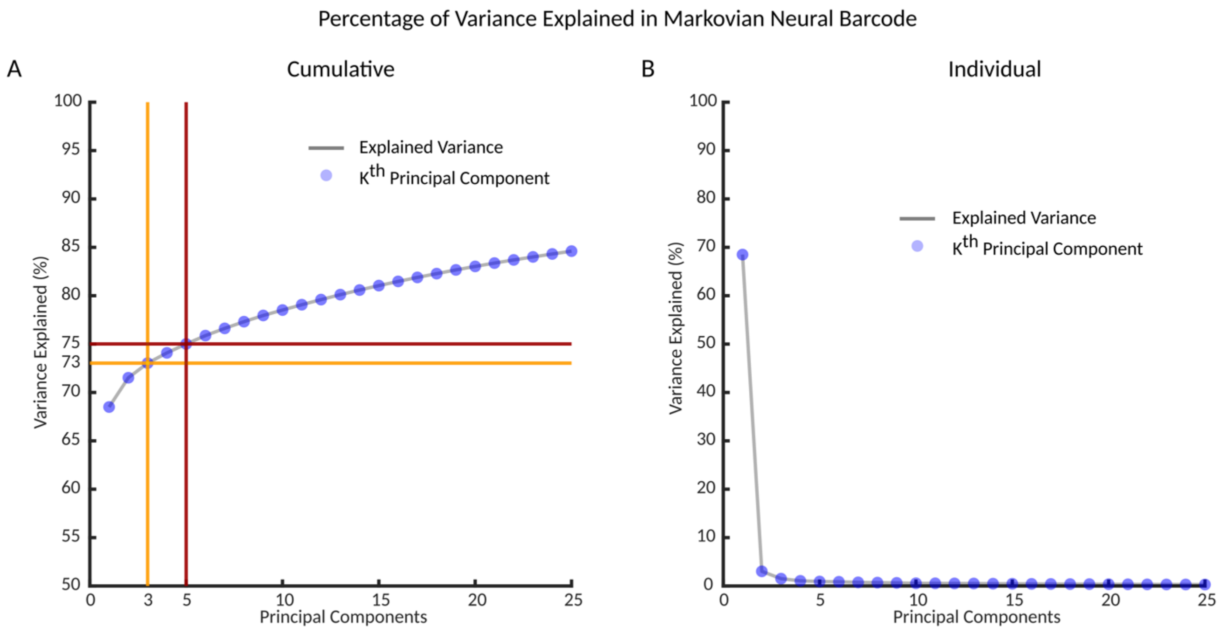


(A) The cumulative percentage of variance explained for the principal component projection of the Markovian neural barcode from $n = 160$ recordings. The top 3 principal components explain 73% of the total variance (orange lines) while the top 5 principal components explain 75% of the variance.

(B) The percentage of variance explained by the $k$th principal component. Notice the sharp drop off after a few principal components, indicating the low-dimensional nature of the Markovian neural barcode.

**Supplemental Figure 6. The Markovian neural barcode is amenable to dimension reduction.**

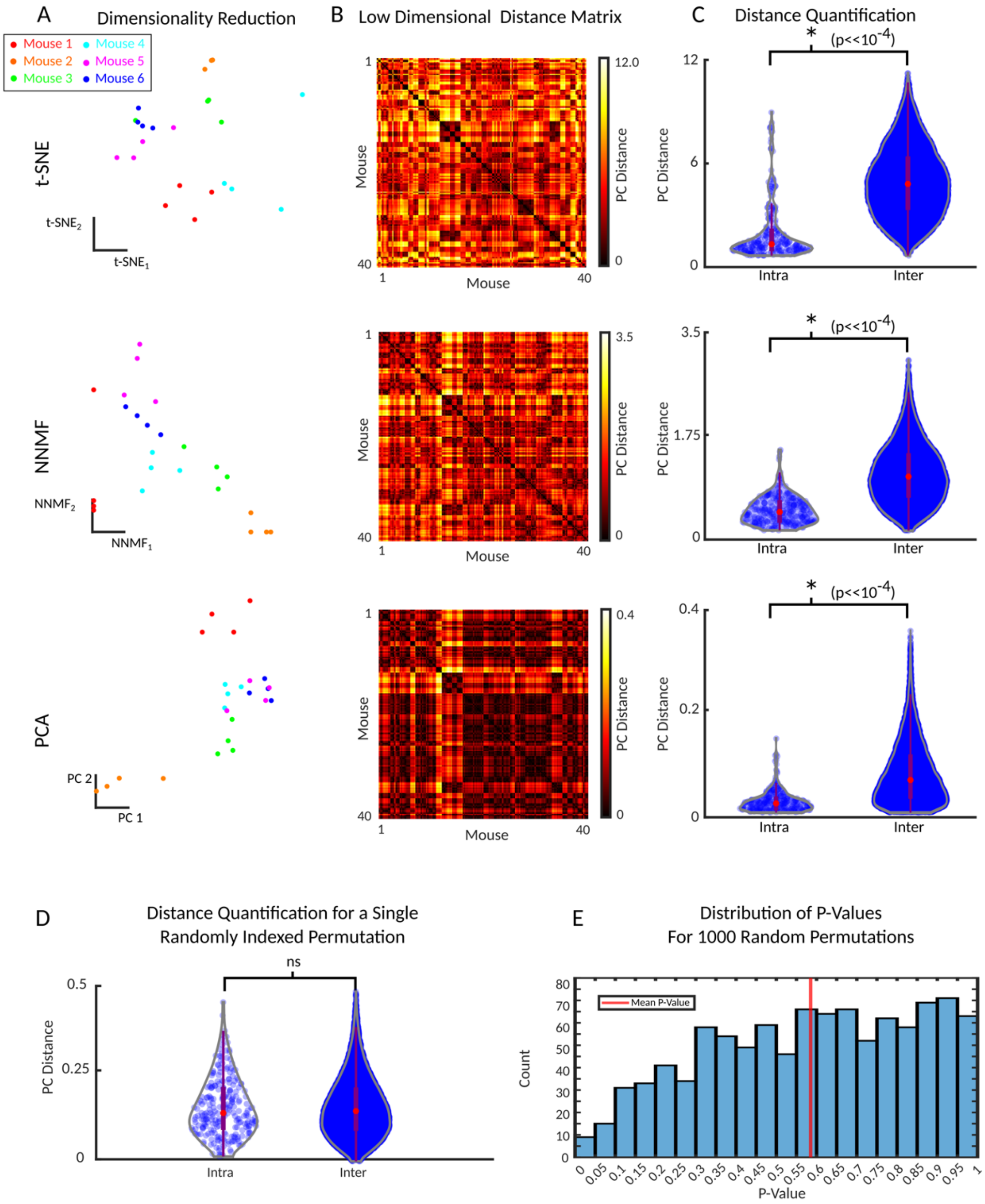

(A) Low-dimensional projections of the neural barcode for the repeated acquisition dataset consisting of 4 recordings (baseline, +1 hour minutes, + 2 hours, + 24 hours) with $n = 40$ mice $(160$ recordings). The dimensions of the Markovian neural barcode were reduced with t-SNE (top), NNMF (middle) and PCA (bottom).

(B) The low-dimensional distance matrix formed by computing the distance between the first 2 components of respective low-dimensional projections in (A). Note the block structure present in all projections, with the dark intra-mouse blocks along the main diagonal.

(C) The intra-mouse distance versus the inter-mouse distance violin plots for t-SNE distances (Top), NNMF distances (Middle), and PCA distances (Bottom). The differences between distances distributions in all cases was significant (Wilcoxon Rank Sum; (t-sne) Intra: $0.79622$, Inter: $4.8433$, $p = 4.3253e - 94$; (NNMF) Intra: $0.36827$, Inter: $1.0504$, $p = 1.0308e - 86$; (PCA) Intra: $0.021259$, Inter: $0.074505$, $p = 5.7549e - 65$). Please see figure for inter-quartile range. The red points in the center of the distance distributions denote the median, while the wider purple lines denote the 25/75 percentile, and the thinner line denote the most extreme points not considered outliers.

(D) The animal-index shuffled intra-animal versus inter-animal distance distributions. The animal indices are randomly shuffled for all $n = 160$ recordings. Distance (PCA) quantification for a single randomly indexed exemplar is given (Wilcoxon Rank Sum test; median: Intra $0.1304$, Inter $0.1292$, $p = 0.9173$). Please see figure for inter-quartile range.

(E) A histogram of the p-values of the Wilcoxon Rank Sum test (mean $\pm$std p-val: $(0.5950 \pm 0.2468)$ in (D) for all 1000 permutations. The probability of a false positive result for the PC-distance blocks is less than 0.01.

**Supplemental Figure 7. The Markovian neural barcode is robust to different sampling rates.**

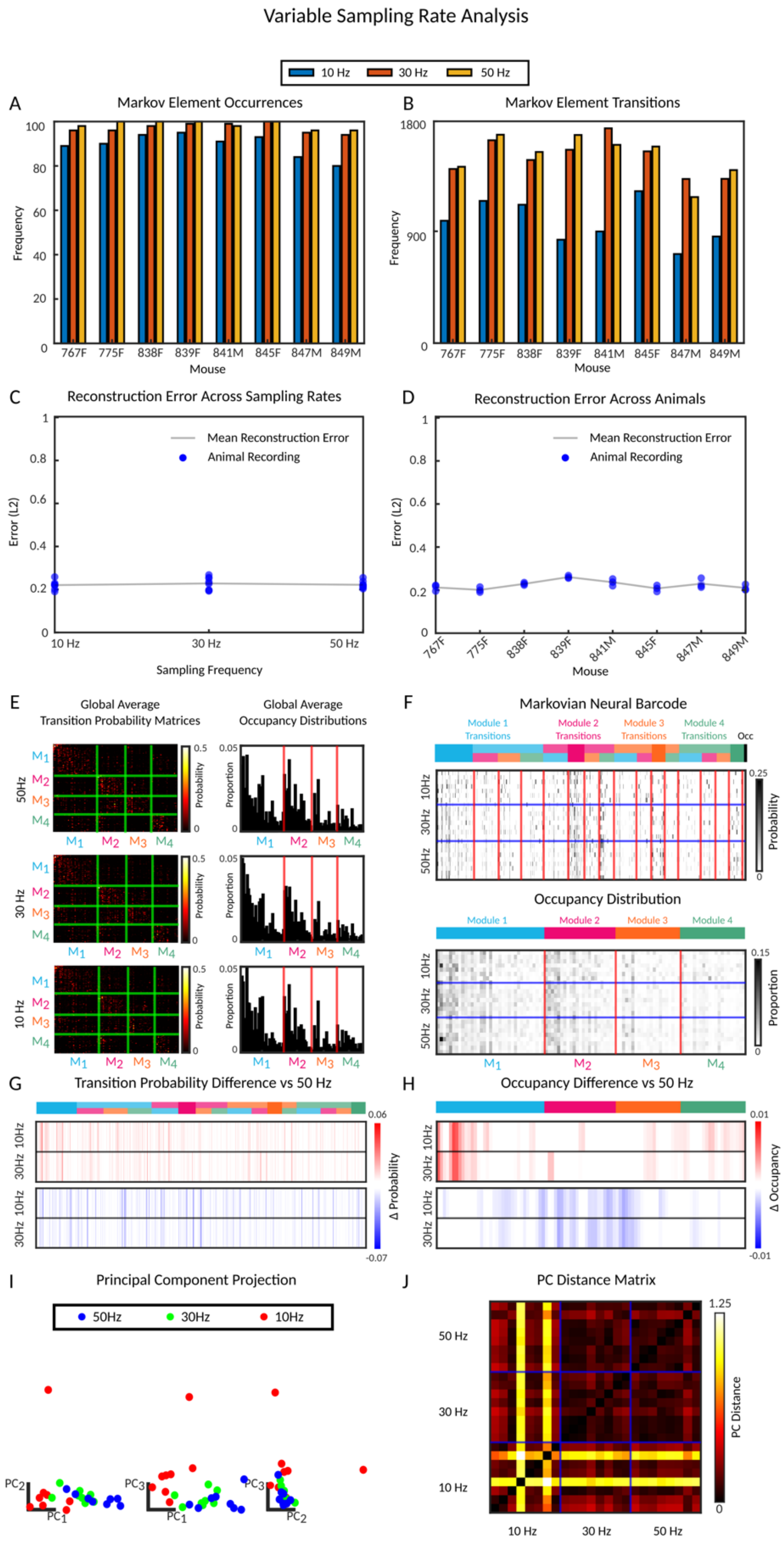

**(A)** A 10Hz acquisition rate resulted in fewer total Markov Element occurrences in recordings matched for acquisition durations, whereas 30Hz and 50Hz were comparable.

**(B)** A 10Hz acquisition rate resulted in fewer Markov Element transitions in recordings matched for acquisition durations, whereas 30Hz and 50Hz were comparable.

**(C)** Imaging acquisition rates did not significantly impact the reconstruction error in a non-overlapping group of $n = 8$ mice.

**(D)** Imaging acquisition rates also did not result in different reconstruction error estimates within the same mouse.

**(E)** Representative transition probability matrices and occupancy distributions are illustrated for the mesoscale cortical activity averaged at 10Hz, 30Hz and 50Hz. Across sampling rates, we can see that there is preservation of the transition probability matrices and occupancy distributions.

**(F)** The Markovian neural barcode derived from $n = 8$ mice imaged at 10Hz, 30Hz, and 50Hz reveals conserved banding and a presented structure across acquisition rates.

**(G)** To visualize the effects of the imaging acquisition rates on the Markovian neural barcode, we illustrate the mean difference between baseline 50Hz and 10Hz and 30Hz acquisition rates to reveal up and downregulated Markov element transitions.

**(H)** The mean difference between 50Hz acquisition and the 10Hz, 30Hz acquisition reveals up and downregulated Markov element occupancies.

**(I)** A low-dimensional representation of the Markovian neural barcode using the first three principal components nevertheless demonstrates clear clustering within the different sampling rates in principal component space.

**(J)** A principal component distance projection facilitated visualization of the differences and similarities in dynamics as shown in **(G)**. Here, the similarity between the 30 and 50 Hz recordings are easily seen in contrast to those at 10 Hz.

**Supplemental Figure 8. Haemodynamic correction of jRGECO1a does not alter the Markovian neural barcode.**

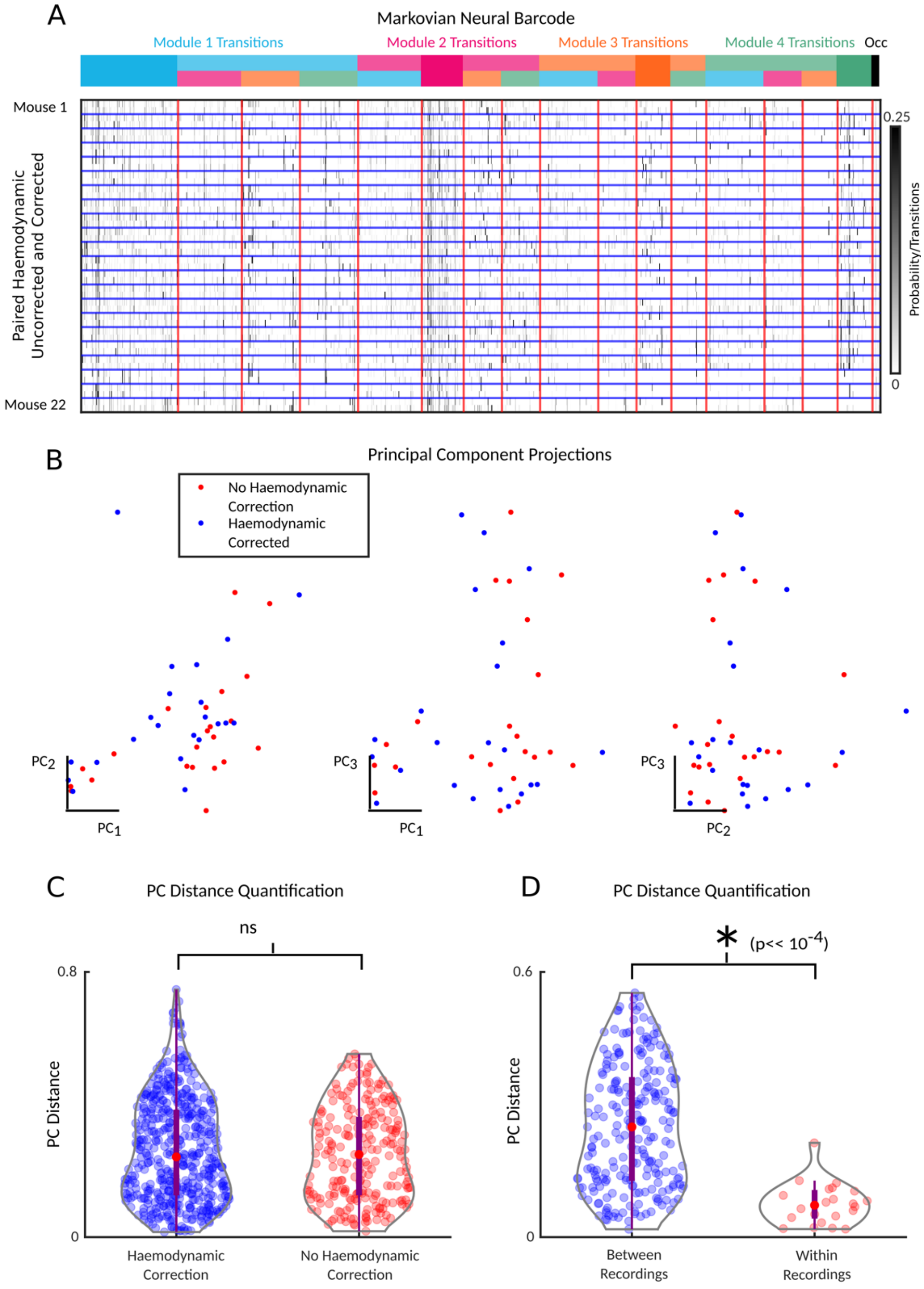

A) The Markovian neural barcode. (Top) A schematic color-coded legend to facilitate visualization and interpretation of the barcode, consisting of the unwrapped Louvain sorted transition probability matrix. The top row of the legend denotes the module of origin while the bottom row of the legend denotes color coded module to which the respective Markov elements transition, followed by the occupancy matrix in black. Each entry corresponds to the barcode derived for the same recording as haemodynamically uncorrected and corrected jRGECO1a activity for n=22 mice.

B) Principal component projection of the Markovian neural barcode derived from jRGECO1a recording without (red) and with (blue) haemodynamic correction applied qualitatively reveals no evidence of clustering.

C) Statistical quantification confirms the absence of a difference between the Markovian neural barcodes of haemodynamically uncorrected and corrected recordings. (Wilcoxon Rank Sum test; Haemo: 0.2421, no-Haemo: 0.2494, $p = 0.5807$)

D) Haemodynamic correction of the jRGECO1a signal has an effect on the Markovian neural barcode for each individual recording, however this is small compared to the differences between animals. (Wilcoxon Rank Sum test; Within: 0.0722, Between: 0.2494, $p = 1.3116e - 94$).

**Supplemental Figure 9. Modelling continuous dynamics.**

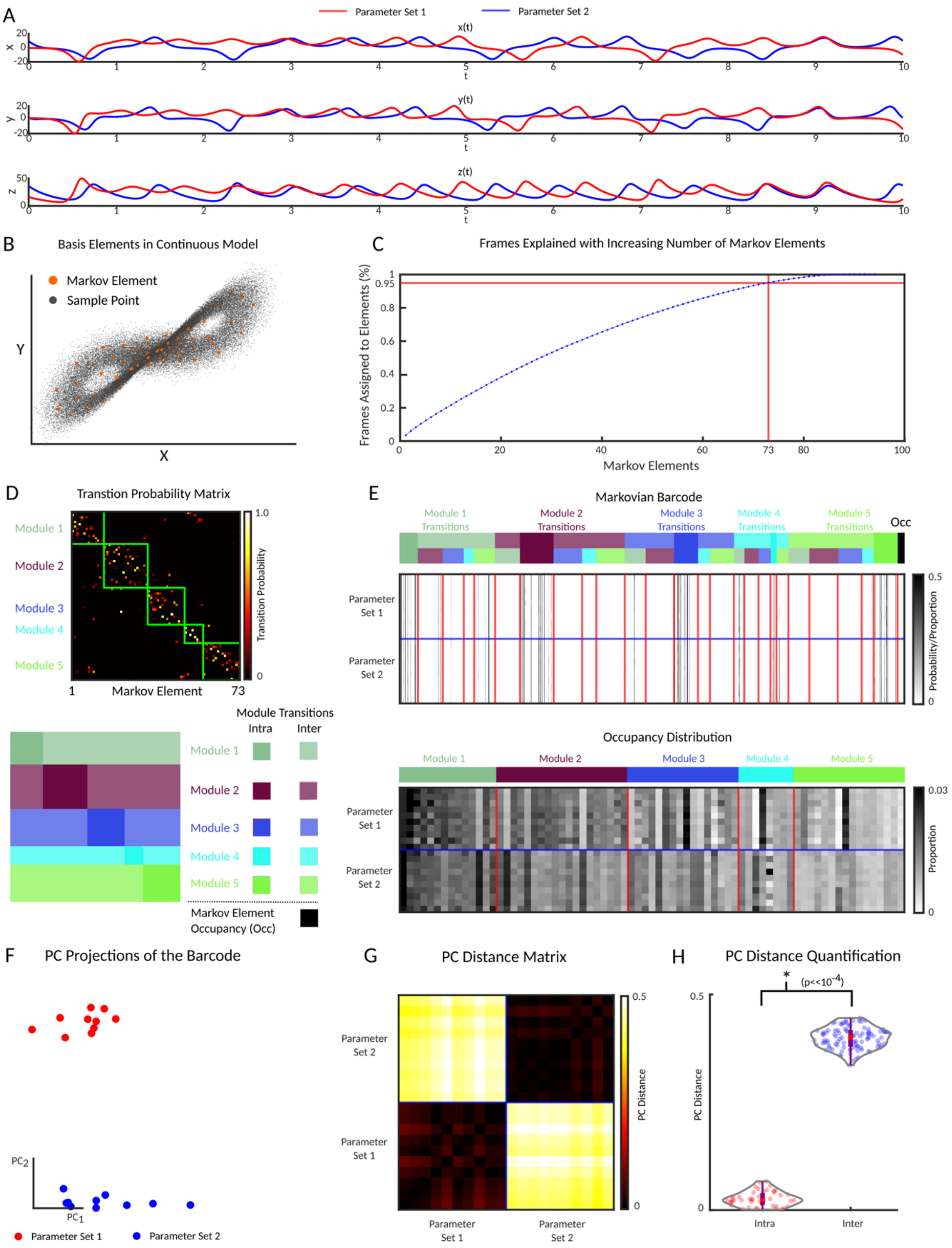

(A) A time series of the Lorenz system for two different parameter sets (red ($\sigma = 10, \beta = 83, \rho = 28,$) and blue ($\sigma = 10, \beta = 83, \rho = 32.$)) . The two trajectories correspond to slightly different chaotic attractors.

(B) The Markov Elements (orange points) constructed from randomly sampling points from both attractors (grey sample points). A total of 73 Markov Elements were selected by SBNMF from 3,000 sample points.

(C) The cumulative distribution of sample points assigned to ranked Markov Elements from SBNMF as a function of increasing size.

(D) (Top) The Louvain sorted transition probability matrix reveals discrete modular dynamics of the Lorenz system. Brighter colours denote larger transition probabilities. (Bottom) The colour legend for Louvain-module to Louvain-module specific transitions.

(E) (Top) The Markovian barcode constructed from the transition probability matrix and the occupancy distribution for the two Lorenz systems (barcodes separated by blue horizontal line). The barcode is delineated by Louvain-module to Louvain-module specific transition blocks. (Bottom) The occupancy distribution component of the Markovian barcode.

(F) The first principal component of the Markovian barcode vs the second principal component for both Lorenz parameter sets (red vs. blue).

(G) The PC distance matrix for parameter for the first two principal components of the Markovian barcode. Note the discrete blocks generated by the slightly different parameter sets and as a result, the slightly different chaotic attractors.

(H) The violin plot (Wilcoxon Rank Sum test; Intra: $0.0286$, Inter: $0.4452$, $p = 6.99e - 22$) of the PC-distance distributions. Please see figure for inter-quartile range. The intra-parameter set 1 distribution $(0.0286)$ is plotted versus the inter-parameter set 2 distribution $(0.4452)$.

**Supplemental Figure 10. Estimating the optimal order of the Markov process.**

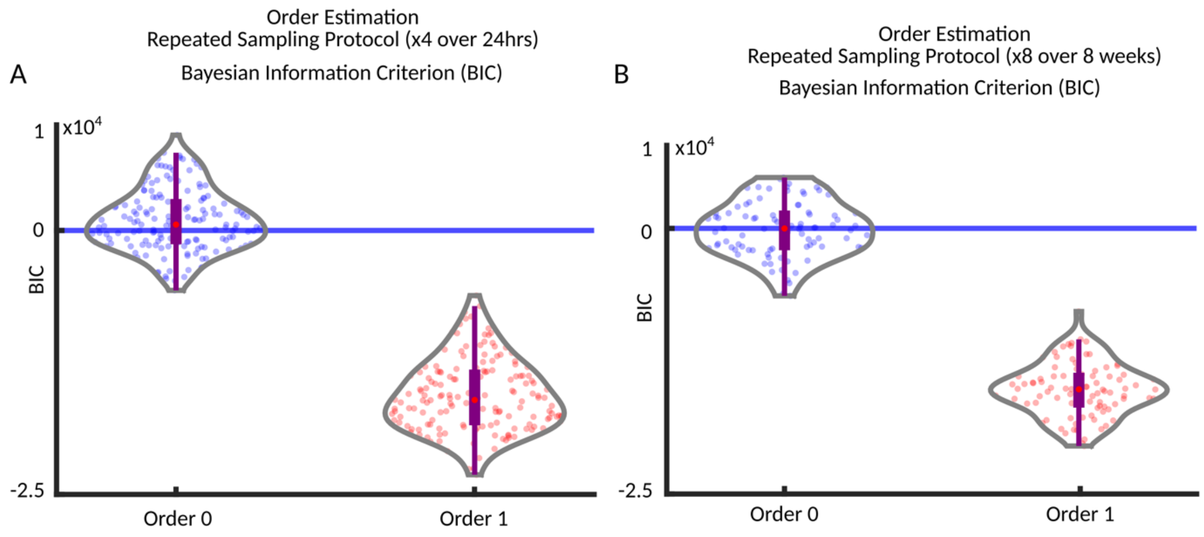


(A) Using Bayesian Information Criterion (BIC) and the $n = 160$ acquisitions, we compared zero and first order Markov models to a second order Markov model. We find that a first order Markov model (median $BIC = -16055.5567$) is always preferred over a second order Markov model, where as a zero order Markov model (median $BIC = 554.4835$) shows no clear preference against a second order model. Please see figure for inter-quartile range.

(B) Using Bayesian Information Criterion (BIC) and an independent $n = 88$ acquisitions, we compared zero and first order Markov models to a second order Markov model. We find that a first order Markov model (median $BIC = -15999.0733$) is always preferred over a second order Markov model, where as a zero order Markov model (median $BIC = 9.2736$) shows no clear preference against a second order model. Please see figure for inter-quartile range.

**Supplemental Figure 11. Convergence of transition probability matrices and the generalizability of the Markovian neural barcode.**

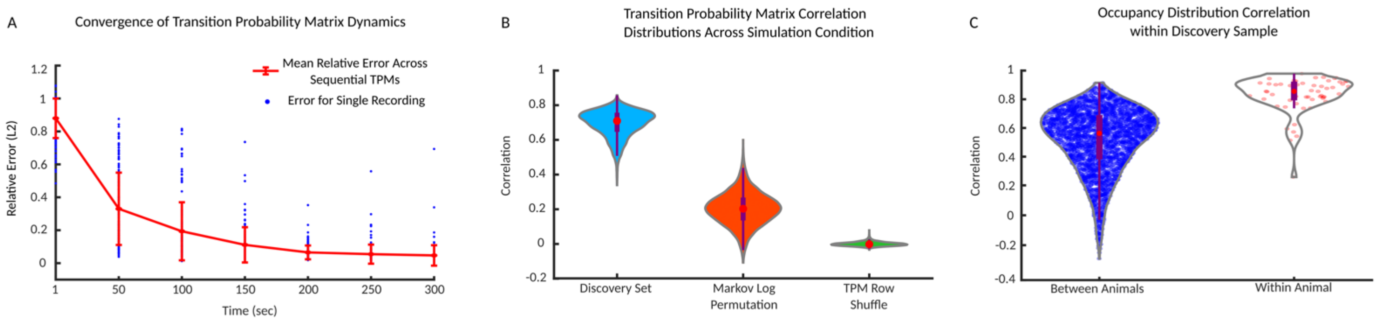


(A) The relative error across sequential transition probability matrices (TPMs) as more frames are used to estimate the temporal dynamics. The relative error is estimated as the matrix 2-norm between TPM computed from Markov logs of increasing temporal size with the full TPM model computed from 15,000 frames. Simulations were computed from the $n = 160$ discovery and validation acquisitions.

(B) The distribution of correlation between the transition probability matrices (TPMs) within recordings from our discovery set ($n = 80$) across different simulation conditions. The median correlation between TPMs in our discovery set was $0.7018$, the median correlation between TPMs generated from permutations of the original Markov logs was lower at $0.1917$, and the median correlation between TPMs generated from row permutations of the original TPMs was $-0.0017$. The median correlation between occupancy distributions was also computed at $0.5684$ across all conditions. Please see figure for inter-quartile range.

(C) To explore the commonality of motifs within and between animals, we computed the correlation between occupancy distributions within and between animals from the $n = 160$ discovery and validation acquisitions. We find that the median correlation within animals was $0.8553$ and median correlation between animals was $0.5623$. These distributions were statistically significant (Wilcoxon Rank Sum Test; Within: $0.8553$, Between: $0.5623$, $p = 1.28e - 27$). Please see figure for inter-quartile range.

**Supplemental Figure 12. Hold-out samples confirm the sensitivity of the Markovian neural barcode to the MES model.**

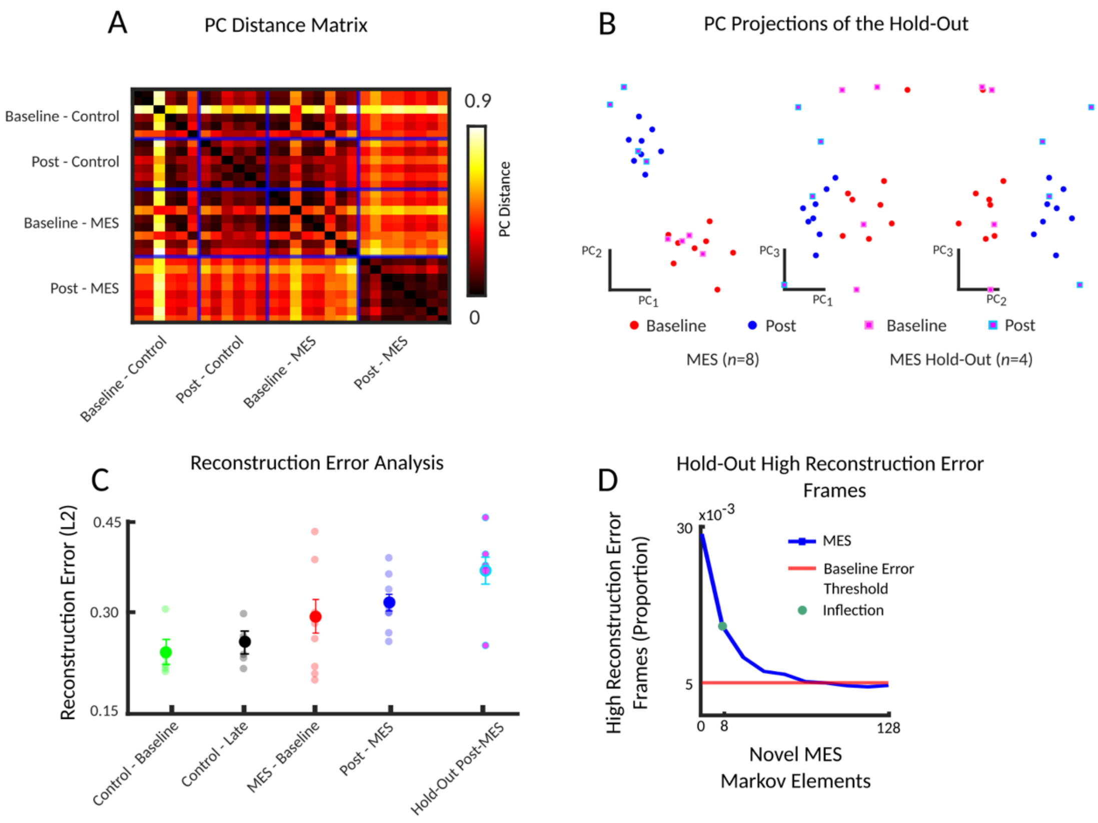


(A) The PC distance matrix for the Maximal Electroconvulsive Seizure (MES) model of seizure and mesoscale cortical imaging in the awake mouse. The blue lines separate mice into different protocols: Baseline (pre MES), post MES, and two time-locked (no MES) controls: Baseline - Control, Post-Control.

(B) The PC-projections of $(n = 8)$ animals under baseline MES (red circle) and POST MES conditions (blue circle), along with $(n = 4)$ hold-out recordings for baseline and POST MES conditions (squares) $(n = 4)$.

(C) The reconstruction error distributions for the Control-Baseline cohort (green), the Control-Late cohort (black), the MES baseline cohort (red), and the Post-MES cohort (blue). The Post-MES holdouts are also plotted (mean $\pm$ se; Control Baseline: $0.2409 \pm 0.0053$, Control Late: $0.2424 \pm 0.0045$, MES Baseline: $0.2753 \pm 0.0085$, Post MES: $0.3012 \pm 0.0040$, HO Post MES: $0.3258 \pm 0.0075$).

(D) The proportion of high reconstruction error frames for the holdouts as more novel, condition specific Markov-Elements derived from the holdout recordings are used.
